# Dissecting the impact of somatic hypermutation on SARS-CoV-2 neutralization and viral escape

**DOI:** 10.1101/2023.05.09.539943

**Authors:** Michael Korenkov, Matthias Zehner, Hadas Cohen-Dvashi, Aliza Borenstein-Katz, Lisa Kottege, Hanna Janicki, Kanika Vanshylla, Timm Weber, Henning Gruell, Manuel Koch, Ron Diskin, Christoph Kreer, Florian Klein

**Author notes:** Contributed equally (Michael Korenkov, Matthias Zehner). Shared senior authorship (Christoph Kreer, Florian Klein).

## Abstract

Somatic hypermutation (SHM) drives affinity maturation and continues over months in SARS-CoV-2 neutralizing antibodies. Yet, several potent SARS-CoV-2 antibodies carry no or only few mutations, leaving the question of how ongoing SHM affects neutralization. Here, we reverted variable region mutations of 92 antibodies and tested their impact on SARS-CoV-2 binding and neutralization. Reverting higher numbers of mutations correlated with decreasing antibody functionality. However, some antibodies, including the public clonotype VH1-58, remained unaffected for Wu01 activity. Moreover, while mutations were dispensable for Wu01-induced VH1-58 antibodies to neutralize Alpha, Beta, and Delta variants, they were critical to neutralize Omicron BA.1/BA.2. Notably, we exploited this knowledge to convert the clinical antibody tixagevimab into a BA.1/BA.2-neutralizer. These findings substantially broaden our understanding of SHM as a mechanism that not only improves antibody responses during affinity maturation, but also counteracts antigenic imprinting through antibody diversification and thus increases the chances of neutralizing viral escape variants.

## INTRODUCTION

B lymphocytes are fundamental for clearing and preventing infections by expressing highly diverse B cell receptors (BCR) that can be secreted as antibodies and target numerous different epitopes. BCR diversity is initially generated by the recombination of variable (V), diversity (D), and joining (J) genes during lymphopoiesis in the bone marrow^1,2^ and further expanded upon antigenic stimuli through somatic hypermutation (SHM) in germinal centers (GC) of secondary lymphoid tissues.^2,3^ Mutations that increase the receptors’ affinity eventually promote the selection and proliferation of high-affinity B cell lineages, underlining the key role of SHM in affinity maturation.^4,5^ More recently, this model has been further refined by revealing not only different fates of cells with varying affinities in the course of the GC reaction but also a strong contribution of stochastic effects that impact repertoire diversity.^6,7^ In line with our current understanding of affinity maturation, reversion of somatic mutations back to germline typically reduces antibody affinity to its cognate antigen and thus biological activity.^8–13^ However, some studies also provide evidence that individual mutations might be neglectable^14–16^ or even detrimental for affinity.^17^

With the emergence of SARS-CoV-2 in late 2019,^18–20^ numerous monoclonal antibodies have been isolated from COVID-19 survivors to gain a deeper understanding of the adaptive humoral SARS-CoV-2 response^21–36^ as well as to treat and prevent infections with therapeutic monoclonal antibodies.^37–43^ The main target of SARS-CoV-2 neutralizing antibodies is the homotrimeric spike (S) protein, which facilitates viral entry into host cells^44–47^ and provides diverse epitopes within the receptor-binding domain (RBD), the N-terminal domain (NTD), and the S2-stem domain.^27,48–50^ Surprisingly, several isolated antibodies were characterized by no or only a low degree of somatic mutations, despite their potent neutralizing activity against the ancestral Wuhan-Hu-1 (Wu01) strain.^30,31,35,36,51^ However, ongoing somatic hypermutation has been demonstrated for SARS-CoV-2 antibodies in the absence of re-exposure,^52–57^ and the requirement of specific mutations for neutralizing activity could be demonstrated for a few selected SARS-CoV-2 antibodies.^58–60^

Despite the success of antibody-mediated treatment and prevention of SARS-CoV-2 infection,^37–43^ emerging SARS-CoV-2 variants with escape mutations in the spike protein have led to reduced vaccine- and infection-induced serum neutralization capacities and rendered approved monoclonal antibodies (mAbs) ineffective *in vitro*.^61–63^ As antibody responses against SARS-CoV-2 can be remarkably convergent, with prominent public clonotypes (e.g., VH3-53/3-66, VH3-30, or VH1-58 classes) developing independently in different individuals,^64–66^ there is an urgent need to investigate how the B cell response copes with viral escape. In particular, understanding the role of somatic mutations is critical to inform on vaccine strategies as well as antibody-based SARS-CoV-2 therapies.

Here, we studied the impact of SHM on SARS-CoV-2 Wu01 spike binding and neutralizing activity in 92 antibodies that were isolated in the first year of the pandemic. Moreover, we selected the highly conserved IGHV1-58/IGKV3-20 public clonotype to decipher the impact of SHM on the neutralizing capacity of emerging viral escape variants and exploited this knowledge to restore the neutralization capacity of clinically-developed antibodies against emerging SARS-CoV-2 variants.

## RESULTS

### The impact of somatic mutations on SARS-CoV-2 Wu01 binding and neutralization diverges for different antibodies

To evaluate the impact of somatic hypermutation on the functionality of SARS-CoV-2 antibodies, we selected 92 monoclonal antibodies from the CoV-AbDab,^67^ which were isolated from at least 33 individuals in 16 distinct studies between 2020 and 2021 and comprise 62 unique VH/VL pairings as well as 86 unique heavy chain complementarity determining region (CDR) 3 sequences (**Figure 1A**, **Supplementary Table 1**). The selected antibodies mainly targeted the receptor-binding domain (RBD, 90/92; **Figure 1B**, **Supplementary Table 1**) and overall carried less mutations than the average memory B cell IgG repertoire^68^ (median of 3.5 [IQR: 2 – 6], 2 [IQR: 0 – 3], and 3 [IQR: 1 – 4] vs 10 [IQR: 6 – 14], 5 [IQR: 3 – 8], [IQR: 3 – 8] amino acid substitutions in heavy, kappa, and lambda chains, respectively; **Figure 1C**). Immunoglobulin heavy, kappa, and lambda chain variable (IGHV, IGKV, and IGLV) gene segments usages reflected the SARS-CoV-2-characteristic overrepresentation of distinct V genes (e.g., IGHV3-53, IGHV3-66, IGKV3-20, and IGLV6-57) with a kappa to lambda ratio of approximately 2:1 (**Supplementary Figure 1**). CDR3 length distributions were comparable to reference memory B cell IgG repertoire data^68^ (**Supplementary Figure 1**). We recombinantly expressed all 92 original wild-type (WT) antibodies as well as the 88 possible VH/VL region germline-reverted (GL) versions (four antibodies with no mutations were used as internal controls) and compared their binding and neutralization activity by Wu01 S-Protein ELISA and pseudo-typed lentivirus neutralization tests. Germline reversion had either no effect or decreased binding and neutralization up to a complete loss of activity (**Figure 1D**, left panel). Moreover, log fold changes in EC_50_ and IC_50_ values between WT and GL variants showed a modest positive correlation with the total number of reverted mutations (**Supplementary Figure 2**), i.e., antibodies with more mutations also more strongly lost functionality upon germline reversion. For instance, antibodies with 6-10 mutations decreased on average 2.4 log_10_ fold in neutralization, while antibodies with 1-5 mutations only decreased by 1.2 log_10_ fold (**Figure 1D**, right panel). We also characterized reactivity against other human coronaviruses (HCoVs) but could not detect any binding against OC43, HKU1, NL63, or 229E spike proteins (**Supplementary Figure 3**). Of note, from the neutralizing antibodies with one or more mutations (88/92), we identified some that were unaffected by germline reversion (no relevant change in EC_50_ and IC_50_; 37/88 and 10/88, respectively, **Figure 1D**). This suggests that these antibodies have only acquired mutations that are mostly neglectable for Wu01 binding and neutralizing activity.

**Figure 1:**
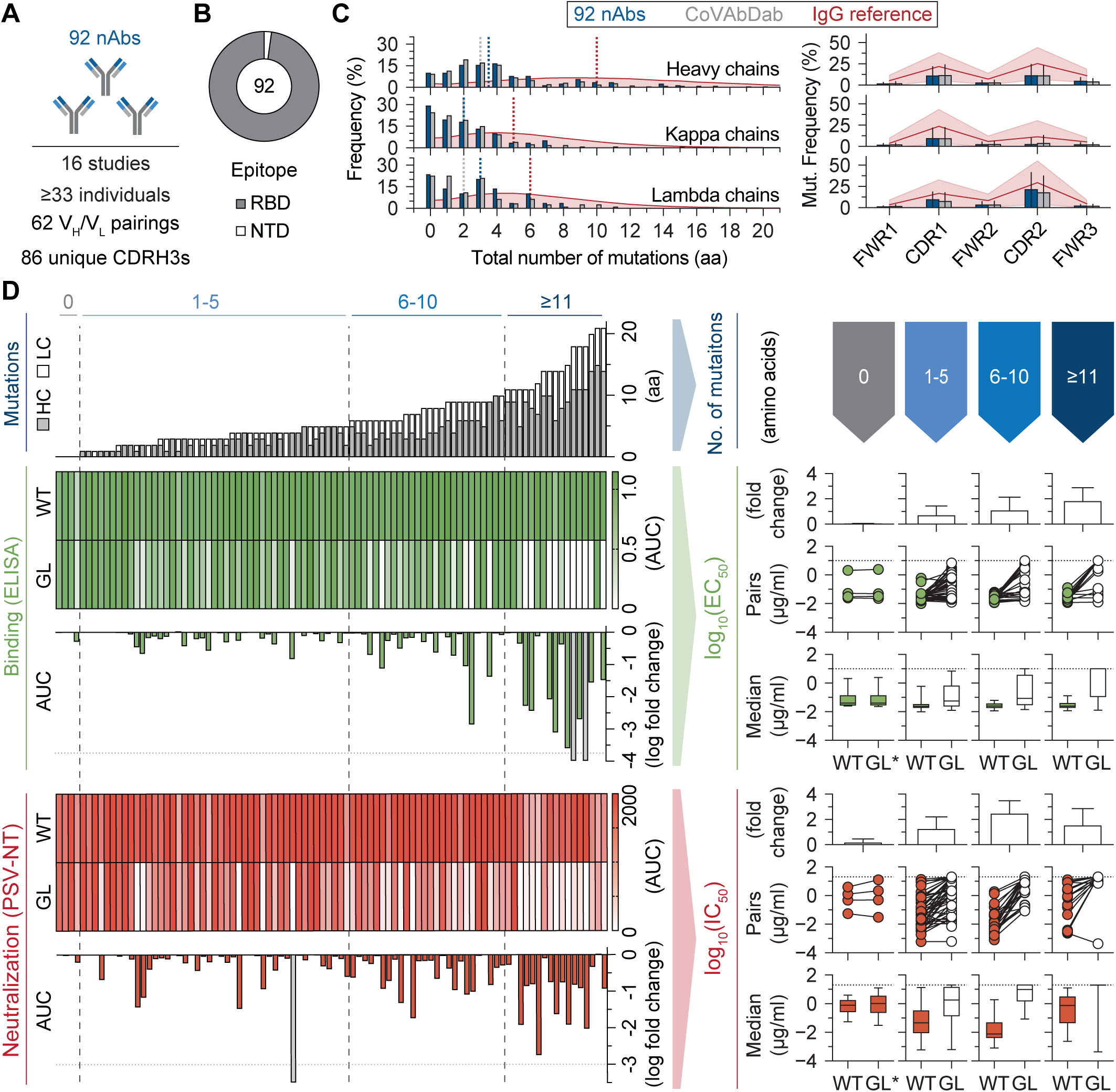
The role of somatic mutations in SARS-CoV-2 Wu01 neutralizing antibodies. (**A**) Composition of SARS-CoV-2 neutralizing antibodies that have been selected from the CoV-AbDab for this study. (**B**) Epitope specificity of the 92 antibodies. (**C**) Distribution of V gene mutations (FWR1 to FWR3) in the selected antibodies in comparison to all human SARS-CoV-2 neutralizing antibodies in the CoV-AbDab at the day of antibody selection, as well as NGS-derived IgG reference sequences of 57 healthy individuals. Left panel shows total number of mutations from FWR1 to FWR3 with dotted lines depicting the median. Right panel shows relative frequency of mutations within FWR and CDR regions. (**D**) Comparison of binding (AUC/EC_50_) and neutralization (AUC/IC_50_) of original wild-type (WT) and the germline-reverted (GL) antibody, after stratifying antibodies by the total number of heavy and light chain amino acid (aa) mutations. Individual log_10_ fold changes in AUC (left panel) and geometric mean log_10_ fold changes in the EC_50_/IC_50_ values (right panel) are given. Fold changes above the dotted lines (open grey bars, left panel) represent complete loss of activity after germline reversion. Upper limits of quantification for EC_50_ and IC_50_ are given as dotted lines (right panel). GL*: For antibodies with no mutations, WT and GL sequences are identical; RBD: receptor binding domain; NTD: N-terminal domain; CDR: complementarity determining region; FWR: framework region; AUC: Area under the curve; PSV-NT: Pseudovirus neutralization test.

We conclude that somatic mutations become increasingly important for potent binding and neutralization of the ancestral Wu01 strain, the more mutations are present in an antibody. However, distinct antibodies are able to act independently of their mutations and rely on either germline V gene or CDR3-encoded sequence features for their activity.

### The public clonotype IGHV1-58/IGKV3-20 neutralizes SARS-CoV-2 Wu01 independently of somatic hypermutation

In order to dissect the role of sequence features that facilitate SHM-independent neutralization of SARS-CoV-2 (i.e., IC_50_ log_10_ fold change ≤0.3 after germline reversion), we investigated the correlation of the IC_50_ log_10_ fold changes with CDRH3 length, hydrophobicity, and V gene segment usage. CDRH3 regions were on average slightly, but not significantly longer as well as more hydrophobic in SHM-independent neutralizing antibodies (**Supplementary Figure 4**). Grouping antibodies by their IGHV gene revealed that the ten antibodies that remained unaffected in neutralization by SHM reversion are distributed across seven different VH/VL gene combinations (**Supplementary Figure 5**). Notably, the majority of common VH gene groups, such as IGHV3-53/3-66, 3-30, and 1-69, dropped substantially in the mean neutralization after germline reversion (**Figure 2A**). Members of the IGHV1-58 group, however, not only ranked among the most potent antibodies, but also acted independently of the few mutations they acquired (**Figure 2A**, highlighted in green). We further investigated this group and found that all tested antibodies belong to a previously described public clonotype (denoted as the VH1-58 class)^64–66^ that is not only marked by the exclusive pairing with the kappa light chain V gene segment IGKV3-20, but also by a highly convergent 16 amino acid CDRH3 region with a double cysteine motif (**Figure 2B**). Notably, as of February 2023, this public clonotype accounts for 95 of the 119 IGHV1-58 antibody sequences in the CoV-AbDab (79%) that are reported to neutralize the ancestral SARS-CoV-2 Wu01 isolate. We and others have solved structures of this public clonotype, revealing contact sites between all CDRs and some residues at the RBD ridge, a long loop that connects β-strands 5 and 6 of the RBD (i.e., residues 473-489) (PDB: 7B0B, **Supplementary Figure 6**, **Supplementary Table 2**).^49,69–71^ Although several RBD residues contribute to the epitope, Phe486 is likely the central one as it is tightly packed in a hydrophobic pocket that is formed at the interface between the heavy and light chains (**Supplementary Figure 6**). Also, a disulfide bridge is forming within the CDRH3, critically contributing to the activity of the antibodies (**Supplementary Figure 6**). Consistent with the sequence similarity, the binding mechanism is highly convergent within the public clonotype, as visualized by the almost identical overlay of seven available structures from VH1-58 antibodies (**Figure 2C**).

**Figure 2:**
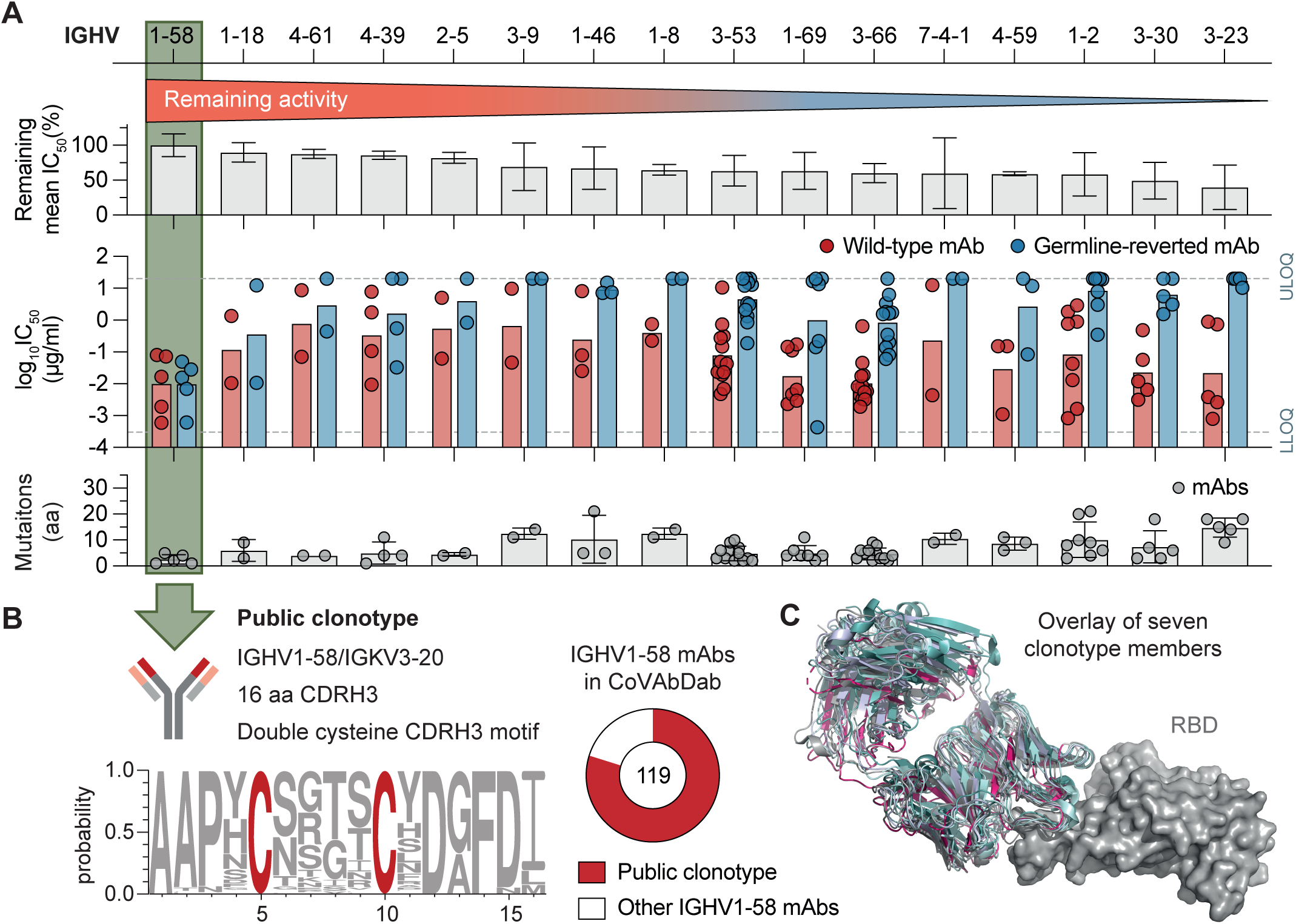
The effect of SHM on neutralization potency within VH gene groups. (**A**) Monoclonal antibodies were grouped by VH genes and ordered according to the mean log_10_ fold IC_50_ change in each group. Top bars depict the mean remaining IC_50_ after germline reversion normalized to the maximum detectable difference, i.e., LLOQ (0.0003 µg/ml) to ULOQ (20 µg/ml). Middle bar graphs represent mean log_10_IC_50_ of wild-type (red) and germline-reverted (blue) antibodies in each group with individual antibodies depicted as dots. Bottom bar graphs show the mean number of mutations per group with individual antibodies depicted as dots. Only groups with at least 2 members are displayed. (**B**) Characteristics of the IGHV1-58/IGKV3-20 public clonotype (left panel) and number of IGHV1-58/IGKV3-20 public clonotype antibodies in the CoV-AbDab (right panel). (**C**) Structural overlay of seven IGHV1-58/IGKV3-20 antibodies binding the SARS-CoV-2 RBD (PDB: 7B0B, 7E3K, 7E3L, 7BEN, 7P40, 7EZV, 7LRS). ULOQ: upper limit of quantification, LLOQ: lower limit of quantification; CDRH3: Heavy chain complementarity determining region 3.

We conclude that antibodies that do not require SHM for potent SARS-CoV-2 neutralization can in principle develop from different V(D)J recombination events as well as heavy and light chain pairings, with the largely SHM-independent VH1-58 public clonotype showing exceptional convergence across many different individuals.

### VH1-58 class antibodies acquired somatic mutations that are required for Omicron BA.1 and BA.2 neutralization

Since VH1-58 antibodies were readily isolated from the first SARS-CoV-2 convalescent individuals, we asked how they cope with viral evolution and what role SHM plays in this context. To this end, we produced another 17 members (22 in total, **Supplementary Table 3**) of the VH1-58 public clonotype and investigated their neutralization against Wu01, Alpha, Beta, Delta, as well as Omicron variants BA.1 and BA.2 in pseudovirus neutralization assays. While all VH1-58 antibodies neutralized Alpha, Beta, and Delta variants, only a subset was able to neutralize the more distant Omicron subvariants (**Figure 3A**). Notably, germline-reversion had no or only minor effects on Wu01, Alpha, Beta, and Delta neutralization, but substantially weakened or completely abolished Omicron neutralization capacity for most Omicron-neutralizing mAbs. This suggests that several VH1-58 members acquired mutations that compensate for differences in the antibody interactions with the spike proteins of earlier variants and that of Omicron. A similar SHM-dependent separation of antibodies by Wu01, Delta, and Omicron BA.1 reactivity was also observed for spike binding (**Supplementary Figure S7**). To find out how mutations promote Omicron neutralization, we investigated the number and pattern of mutations in the 22 VH1-58 antibodies. Overall, somatic mutations are enriched in heavy chains of the Omicron-neutralizing antibody subset with potential mutational hotspot regions and shared mutations (e.g., T30S in CDRH1, **Figure 3B**). Light chain mutations seem to be focused on CDRL1 and 2, with a prominent S30R mutation in the CDRL1 (**Figure 3B**). Notably, heavy chains from Omicron neutralizing antibodies mostly retained neutralizing capacity when paired with light chains from non-Omicron neutralizing antibodies, but not vice versa, suggesting that heavy chain mutations more strongly contribute to Omicron neutralization (**Figure 3C**). To identify critical amino acid substitutions in those heavy chains, we reverted individual or combined heavy chain CDR as well as FWR mutations in all Omicron neutralizing antibodies (**Supplementary Figure 8A**). Moreover, we generated single as well as combinational mutation reversions for the two most potent Omicron neutralizing antibodies C043 and CQTS004 (**Supplementary Figure 8B**). Almost all antibodies tolerated the partial removal of mutations from heavy chain CDRs or FWRs, with neutralization capacities decreasing up to one log_10_ fold, but not vanishing completely. This suggests that these heavy chain mutations mostly act in an additive manner to convey Omicron neutralization. However, for CQTS004, we could identify N47D in the CDR2 as a critical single substitution with a major impact on omicron neutralization. Notably, some reversions in CQTS004 (including N47D) actually increased neutralizing activity against Wu01.

**Figure 3:**
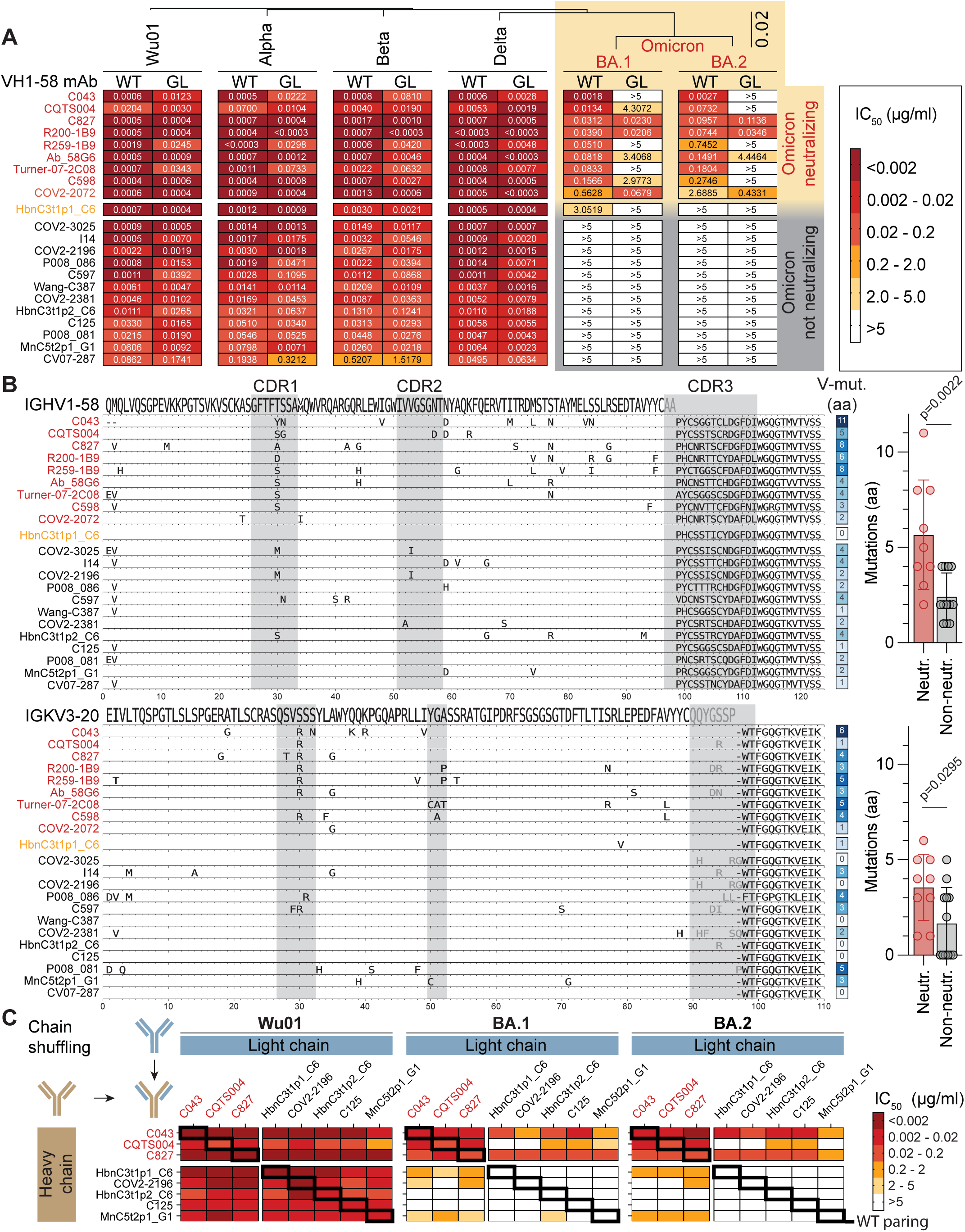
The impact of SHM in VH1-58 antibodies on SARS-CoV-2 variant neutralization. (**A**) 22 original wild-type (WT) and germline-reverted (GL) antibodies were tested for neutralization against Wu01-, Alpha-, Beta-, Delta-, BA.1-, and BA.2-pseudotyped virus. Heatmaps show mean IC_50_ values of at least two independent experiments. Antibodies are ranked by IC_50_ values against BA.1. The phylogenetic tree shows amino acid distances between SARS-Cov-2 Spike variants with the scale bar indicating 0.02 substitutions per site. (**B**) Sequence alignment of IGHV1-58 heavy (top panel) and IGKV3-20 light chains (lower panel) with V gene amino acid mutations (FWR1 to FWR3, excluding the CDR3). Bar graphs depict amino acid mutation means of (n=9) neutralizing vs. (n=12) non-neutralizing antibodies, excluding HbnC3t1p1_C6. P values were determined by unpaired two-tailed t tests. (**C**) Shuffled heavy (rows) and light chains (columns) from VH1-58 antibodies were tested for neutralization against Wu01, BA.1, and BA.2. Heatmaps show mean IC_50_ values of two technical replicates. aa: amino acid; CDR: complementarity determining region.

We conclude that some VH1-58 antibodies elicited by pre-Omicron SARS-CoV2 variants have accumulated different “bystander” mutations that additively convey neutralizing activity against Omicron BA.1 and BA.2.

### Identification of single mutations that confer Omicron BA.1 and BA.2 neutralizing activity on non-neutralizing VH1-58 antibodies

COV2-2196 (tixagevimab) is a VH1-58 antibody that has been used in combination with the antibody cilgavimab for treatment and pre-exposure prophylaxis of SARS-CoV-2 infection, but is ineffective against Omicron BA.1 and BA.2.^61^ Since CDRH3s of the VH1-58 clonotype are highly convergent, we speculated that VH gene mutations are interchangeable between clonotype members.

We therefore transferred the fully mutated VH genes from eight Omicron neutralizing antibodies (donor) onto six non-as well as weakly Omicron neutralizing mAbs (acceptor), retaining the acceptors light chains as well as their original CDRH3s. Strikingly, we could fully restore BA.1 and BA.2 neutralizing activity in all non-neutralizing antibodies with at least four different VH gene mutation patterns and concordantly improve the neutralizing capacity of weak neutralizers (**Figure 4A**). Interestingly, even though the mutations from antibodies R200-1B9 and C827 were dispensable for their own Omicron-neutralizing activity (**Figure 3A**), they still can confer Omicron neutralization on other VH1-58 antibodies. We then selected ten mutation patterns with single as well as combinations of up to five amino acid substitutions from the Omicron neutralizing antibodies and introduced them in the heavy chains of three partially or non-Omicron-neutralizing antibodies, including COV2-2196 (**Figure 4B**). For HbnC3t1p1_C6, all ten mutation patterns improved BA.1, and nine out of ten patterns restored BA.2 neutralization. For COV2-2196, seven out of ten combinations restored both BA.1 and BA.2 neutralization, whereas for C125, seven out of ten combinations restored BA.1, and four out of ten combinations restored BA.2 neutralization. The different degrees of improved neutralization might be explained by the initial capacity of the antibodies to bind Omicron. Although WT HbnC3t1p1_C6 did merely neutralize BA.1 and failed to neutralize BA.2 within the range of concentrations tested, it had the lowest EC_50_ against BA.1 spike (0.5 µg/ml) out of the three antibodies, followed by COV2-2196 (1.6 µg/ml) and no detectable binding for C125 (>10 µg/ml, **Supplementary Figure 7**). In line with this, the CDRH3 of HbnC3t1p1_C6 is also more similar to the SHM-independent Omicron neutralizers R200-1B9, C827, and COV2-2072 with a histidine at position 100 and an aromatic residue (tyrosine) at position 107. Importantly, HbnC3t1p1_C6 as well as COV2-2196 could be modified to potently neutralize Omicron BA.1 and BA.2 by inserting a single aspartic acid at position 57, which is the same substitution that turned out to be critical for Omicron neutralizing activity in CQTS004 (**Supplementary Figure 8B**). In the BA.1/BA.2 spike proteins, an arginine residue replaces glutamine in position 493, abrogating a strong double hydrogen bond interaction with Asn57 of HbnC3t1p1_C6. Hence, introducing aspartic acid in position 57 could restore the ability of the antibody to target the RBD by enabling the formation of a salt bridge between Asp57 and Arg493.

**Figure 4:**
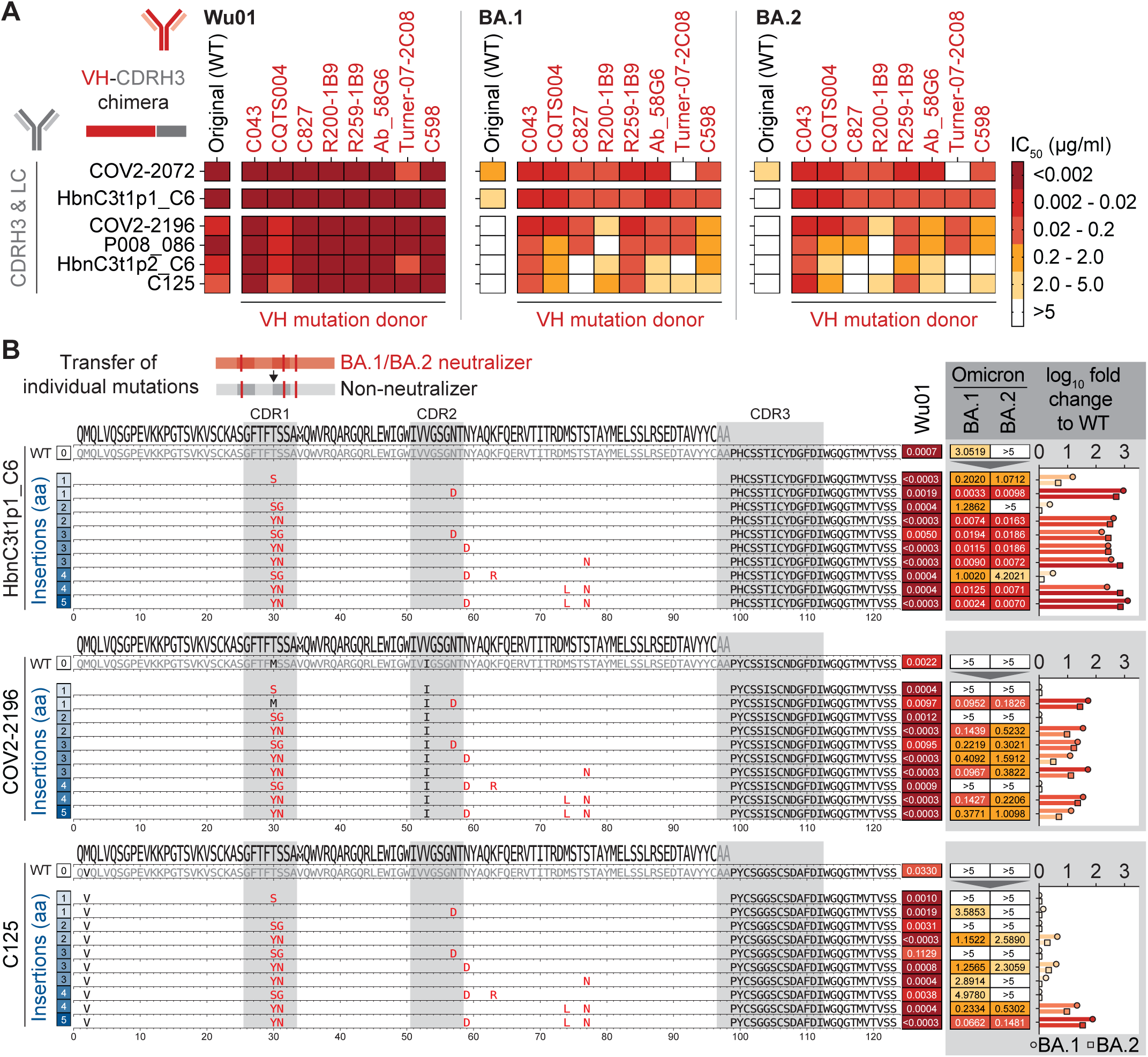
SHM transfer from Omicron neutralizing to non-neutralizing VH1-58 antibodies. (**A**) VH1-58 antibody VH-CDRH3 chimeras were generated by replacing VH regions (FWR1 to FWR3) of BA.1/2 weakly, partially, or non-neutralizing VH1-58 antibodies with potently BA.1/2 neutralizing antibody VH regions, while retaining the original CDRH3 and complete light chain (LC). CDRH3/LC donors are depicted on the left as rows, VH mutation donors are depicted on the top as columns. Heatmaps show Wu01, BA.1, and BA.2 mean IC_50_ values of two independent experiments for VH-CDRH3 chimeras. IC_50_ values of original wild-type antibodies were taken from Figure 3A for comparison. (**B**) Amino acid sequence alignment of HbnC3t1p1_C6, COV2-2196, and C125 wild-type antibodies with IGHV1-58, as well as individual mutation variants ordered by the number of inserted mutations. Heatmaps to the right show mean IC_50_ values of two independent experiments against Wu01, BA.1, and BA.2. Bar graphs depict log_10_ fold change IC_50_ against BA.1 (circles) and BA.2 (squares) to original wild-type antibodies with values “>5” set to 5 for calculations. WT: wild-type; CDR: complementarity determining region; aa: amino acid.

We conclude that a single amino acid substitution is sufficient to restore Omicron neutralizing activity in VH1-58 antibodies, including the clinical antibody tixagevimab, which can have major implications to more rapidly adapt and develop therapeutic monoclonal antibodies to target novel SARS-CoV-2 variants.

## DISCUSSION

SARS-CoV-2 neutralizing antibodies, which are key to the protective immune response upon natural infection or vaccination, developed highly convergent across the population with predominant public clonotypes (e.g. VH1-58, VH3-30, VH3-53/3-66).^64–66^ Notably, numerous potent neutralizing antibodies showed consistently low degrees of SHM,^30,31,35,51,72^ while subsequent studies demonstrated a continuous accumulation of mutations, even in the absence of re-infection or booster vaccination.^52–57^ It thus remained elusive to which degree SHM and germline-encoded features contribute to SARS-CoV-2 binding and neutralization.

By reverting heavy and light chain V gene mutations in 92 monoclonal SARS-CoV-2 Wu01-neutralizing antibodies, we show that mutations in these antibodies mostly improved binding and neutralization, which is in line with previous studies on selected antibodies targeting SARS-CoV-2 or other viruses.^59,73–76^ In addition, we found a moderate positive correlation between the drop of antibody activity and the total number of reverted mutations, implying that antibodies with more mutations also more strongly depend on them. We considered that these antibodies could have developed from cross-reactive HCoV-induced memory B-cells, as described previously.^77,78^ However, none of the wild-type or germline variant antibodies showed any reactivity against OC43, HKU1, NL63, or 229E spike proteins, supporting previous studies that found only little or no recall from memory HCoV responses.^79,80^

In the context of other pathogens including HIV-1, HCV, or RSV, it has been shown that antibodies carry mutations that are neglectable for binding or neutralization.^14–16^ Concordantly, we identified a fraction of mutated anti-SARS-CoV-2 antibodies (∼11%) that bound and neutralized Wu01 independently of acquired mutations, at least within the resolution of our assays. While most of these antibodies were unrelated by sequence, a subgroup consisted of the highly convergent VH1-58 public clonotype class. Mutations that were neglectable for Wu01-reactivity in these antibodies might have been selected against subliminal viral mutants, to eliminate auto reactivity,^3,81^ or because of surface expression effects that impact clonal selection.^3^ However, we propose that they were passing affinity maturation as “bystander” mutations with only little impact on selection processes. The propagation of such mutations could also be the result of the recently described random “clonal bursts” in germinal center reactions that partially uncouple SHM and affinity maturation.^5^

With the ongoing emergence of increasingly mutated escape variants, protection through Wu01- and vaccine-induced serum antibodies as well as approved monoclonal antibodies can be significantly altered.^61–63,82^ Here, the abundant public clonotype VH1-58 with its highly convergent CDRH3 motif and binding mode can serve as a prototype of an imprinted antibody response to investigate the isolated impact of pre-existing mutations on viral escape. In our study, all VH1-58 members were isolated prior to the emergence of Omicron and were able to neutralize SARS-CoV-2 variants Wu01, Alpha, Beta, and Delta independently of SHM. However, only a subset showed Omicron neutralizing activity, which was substantially impaired or lost upon germline reversion, demonstrating the requirement for mutations that have been acquired before the occurrence of Omicron. Biologically, this finding aligns with the observation that antibodies from late B cell lineage members neutralized escape mutations that were emerging in vitro in the presence of the corresponding early predecessor antibodies.^57^ Notably, we observed different mutation patterns that restored Omicron neutralizing activity in VH1-58 clonotype members, suggesting that a divergent evolution of the otherwise convergent clonotype took place in different hosts. In this sense, diversification through ongoing SHM can be beneficial for the immune system to cope with antigenically drifted pathogens in the future. In particular, while some somatic mutations have no substantial impact on effectively targeting the initial pathogen, they play a critical role in retaining repertoire diversity and flexibility that countermeasures antigenic imprinting.

By transferring “bystander” mutations from potent BA.1/BA.2 neutralizing VH1-58/VK3-20 antibodies to non-neutralizing clonotype members, we were able to induce Omicron neutralization in the latter. Moreover, a single substitution was sufficient to restore the neutralizing activity of the parental tixagevimab antibody COV2-2196, which provides a proof of concept that therapeutically-applied antibodies can be minimally adapted to overcome escape mechanisms posed by viral variants.

In conclusion, we present evidence that an appreciable number of SARS-CoV-2 neutralizing antibodies contain mutations with only little contribution to the selection against the ancestral Wu01 strain. However, these “bystander” mutations are beneficial to cope with upcoming variants by increasing the diversity of the imprinted pool of memory B cells.

## Supporting information

Supplementary Information Korenkov et al.

## ACKNOWLEDGEMENTS

We thank all members of the Klein Laboratories for continuous support and helpful discussion; This work was funded by grants from the German Center for Infection Research (DZIF to M.K. and F.K.), the German Research Foundation (DFG; CRC 1279, F.K.; CRC 1310, F.K. and C.K.), the European Research Council (ERC-StG639961, F.K).

## AUTHOR CONTRIBUTIONS

Conceptualization, F.K, C.K., M.Z.; Methodology, F.K., C.K.; Investigation, M.K., M.Z., H.C., K.V., H.G., A.B. L.K., H.J., Ma.K., T.W.; Software, C.K., M.K.; Formal Analysis, M.K., H.C., M.Z., R.D., and C.K.; Resources, K.V., H.G., Ma.K., R.D., F.K.; Data Curation: M.K., C.K.; Writing - original Draft, M.K., C.K.; Writing - review and editing, all authors; Visualization - M.K., C.K., R.D.; Supervision, R.D., C.K., F.K.; Funding acquisition - C.K., F.K.

## SUPPLEMENTARY FIGURE LEGENDS

**Supplementary Figure 1: Sequence features of the 92 selected antibodies.** (**A**) Heavy chain V gene segment usage and complementarity determining region (CDR) 3 length distributions from the selected 92 antibodies in comparison to the complete 319 human SARS-CoV-2 neutralizing antibodies from the CoV-AbDab and IgG reference repertoires from n=57 healthy individuals. (**B**) Kappa and lambda fraction of light chains from the 92 selected antibodies and the CoV-AbDab antibodies, as well as V gene segment usage and CDR3 length distributions from the selected antibodies, the CoV-AbDab antibodies, and IgG reference repertoires.

**Supplementary Figure 2:** Correlation of change in binding or neutralization after germline reversion with the number of reverted mutations. Log_10_ fold changes in EC_50_ (binding) or IC_50_ (neutralization) after germline-reversion are plotted against the total number of reverted heavy and light chain mutations. Spearman correlation coefficients r_s_ as well as corresponding p values are depicted in the plots. aa: amino acid.

**Supplementary Figure 3: Cross-reactivity of wild-type and germline-reverted antibodies with endemic human coronavirus spike proteins.** Area under the curve (AUC) heatmap of ELISAs against Wu01, OC43, HKU1, 229E, and NL63 spike proteins. Human coronaviruses are ordered by their phylogenetic distance on amino acid (aa) level. The tree scale represents one amino acid substitution per site. Five control sera of healthy individuals were taken as positive controls for human coronavirus spike proteins. The upper part including the mutation count and Wu01 AUC data for individual wild-type and germline-reverted antibodies was taken from Figure 1D for comparison.

**Supplementary Figure 4: Correlation of change in neutralization after germline reversion with CDRH3 length and hydrophobicity.** Log_10_ fold change IC_50_ (neutralization) after germline-reversion is plotted against the CDRH3 length (aa: amino acids) and hydrophobicity (based on the Eisenberg scale) for antibodies that contained at least one mutation and did not lose neutralizing activity completely after germline reversion (n=58). Boundary for antibodies to count as unaffected by reversion was set to a log_10_ fold change ≤0.3. Spearman correlation coefficients r_s_ and corresponding p values as well as p values from two-tailed, unpaired t tests are depicted in the plots. CDR: complementarity determining region.

**Supplementary Figure 5: The effect of SHM on binding and neutralization potency grouped by VH and VL genes** Monoclonal antibodies were grouped by VH and VL genes and sorted by the mean log_10_ fold IC_50_ change in each group as in Figure 2. Upper panel: Bars depict the mean remaining EC_50_ after germline reversion normalized to the maximum detectable difference, i.e., lower limit of quantification (LLOQ, 0.0001 μg/ml) to upper limit of quantification (ULOQ, 10 μg/ml). Scatter plots show log_10_ EC_50_ values of individual antibodies before (squares) and after (circles) germline-reversion with corresponding pairs connected by a line. ULOQ is depicted as a dashed line. Antibodies that remained unaffected by reversion (log_10_ fold EC_50_ change ≤ 0.3) are highlighted in green. Middle panel: Bars depict the mean remaining IC_50_ after germline reversion normalized to the maximum detectable difference, i.e., LLOQ (0.0003 μg/ml) to ULOQ (20 μg/ml). Scatter plots show log_10_IC_50_ values of individual antibodies before (squares) and after (circles) germline-reversion with corresponding pairs connected by a line. ULOQ/LLOQ are depicted as dashed lines. Antibodies that remained unaffected by reversion (log_10_ fold IC_50_ change ≤ 0.3) are highlighted in red. Lower panel: Individual (dots) and mean (bars) number of mutations in the VH/VL group. Error bars depict standard deviation, where applicable.

**Supplementary Figure 6: Crystal structure of HbnC3t1p1_C6 and role of the disulfide bridge in the CDRH3** (**A**) The overall structure of the HbnC3t1p1_C6 Fab (blue and green for the heavy and light chains, respectively) in complex with the SARS-CoV-2 RBD (grey, semitransparent surface representation). The epitope of HbnC3t1p1_C6 is highlighted in orange, and Phe486, which is a central feature of the complex, is marked. The greyed inset shows the HbnC3t1p1_C6/RBD complex superimposed with the ACE2/RBD structure (PDB ID: 6M17). HbnC3t1p1_C6 and ACE2 substantially overlap with each other, indicating that neutralization is achieved by blocking the interaction with ACE2. (**B**) An overview of the HbnC3t1p1_C6 epitope. The residues of SARS-CoV-2 RBD that make the epitope of HbnC3t1p1_C6 are labeled and shown as orange sticks. The heavy and light chains of HbnC3t1p1_C6 are shown using a semitransparent surface in blue and green, respectively. The CDRs are labeled. (**C**) HbnC3t1p1_C6 makes a hydrophobic pocket that interacts with Phe486 of the RBD at the interface between its heavy and light chains. Tyr33 from CDRL1, Tyr92, and Trp97 from CDRL3 of the light chain make part of the hydrophobic pocket and are indicated. At the heavy chain, Pro99 at the base and Phe110 at the end of CDRH3 make the bottom of the pocket. The side of the pocket is made of Trp50, a framework residue of the heavy chain. Val52 of CDRH2 makes hydrophobic interaction with Tyr489 of the RBD. (**D**) CDRH3 makes polar interactions with RBD. The main chain of CDRH3 is shown as sticks. Cys106 and Cys101 make a disulfide bridge that, together with a series of hydrogen bonds (green dashed lines) that involve mainchain atoms, stabilize the conformation of CDRH3. Asp108, through its sidechain, and Cys106 and Tyr107, through their mainchain, make hydrogen bonds (yellow dashed lines) with Thr478, Ser477, Asn487, and Ala475 of the RBD. (**E**) CDRH2 makes polar interaction with RBD. Asn57 is stabilized by hydrogen bonds (green dashed lines) to the nearby Ser55 and further forms a hydrogen bond (yellow dashed line) with Gln493 of RBD. (**F**) Binding and neutralization of HbnC3t1p1_C6 with single (SC, CS) and double (SS) cysteine substitutions to serine.

**Supplementary Figure 7: The impact of SHM on binding of VH1-58 antibodies to Wu01, Delta, and Omicron BA.1 spike protein.** EC_50_ values were determined by ELISA against Wu01, Delta, and Omicron BA.1 trimeric spike protein for original wild-type and germline-reverted antibodies. Values represent biological duplicates. Red labels depict Omicron BA.1 and BA.2 neutralizing antibodies.

**Supplementary Figure 8: The impact of grouped and individual mutation reversions on Omicron neutralizing VH1-58 antibodies.** (**A**) Heavy chain CDRs or FWRs were reverted to germline in eight Omicron neutralizing antibodies and partially reverted antibodies (paired with original unreverted light chains) were tested for neutralization against Wu01, BA.1, and BA.2. Heatmaps depict mean IC_50_ values for partially reverted antibodies from two independent experiments. (**B**) Additional individual or combinations of heavy chain mutations were reverted for CQTS004 and C043 as illustrated by blue amino acids in the sequence alignment and IC_50_ values were determined. IC_50_ values represent means of two independent experiments. Reversion variants in (B) are sorted by the number of reverted amino acids. Original wild-type (WT) neutralization values were taken from Figure 3A. CDR: complementarity determining region; FWR: framework region; aa: amino acid.

## METHODS

### RESSOURCES AVAILABILITY

#### Lead Contact

Further information and requests for resources and reagents should be directed to and will be fulfilled by the Lead Contact, Florian Klein.

#### Materials Availability

Requests for materials will be fulfilled by the Lead Contact upon request and might require a Material Transfer Agreement (MTA) for non-commercial usage.

#### Data and Code Availability

All data that support the findings of this study are available within this article. Any additional code to process the data will be shared by the Lead Contact upon request.

### METHOD DETAILS

#### Selection of SARS-CoV-2 neutralizing antibodies

For the initial screening, SARS-CoV-2 neutralizing monoclonal antibodies were selected from previously isolated antibodies^36^ as well as randomly selected from the Coronavirus Antibody Database^67^ (i.e., in total from 319 human B-cell origin and SARS-CoV-2 neutralizing antibodies with complete heavy and light chain sequences by 20.01.2021). Wild-type antibodies were produced and re-evaluated for their neutralizing activity in a Wuhan-Hu1 Spike protein pseudo-typed lentiviral neutralization assay (see below for method details). For inclusion into further analyses, antibodies were required to achieve an IC_50_ of at least 20 µg/ml in the in-house neutralization test, yielding a final set of 92 antibodies. For in-depth analyses of IGHV1-58 public clonotype members 17 additional antibodies have been selected from the CoV-AbDab (retrieved on 09.09.2021) and our own study.^83^

#### Antibody germline reversion

After selection, amino acid sequences were annotated using igblastp (igblast 1.16.0 package)^84^ and reverse translated using the reverse translate tool from the Sequence Manipulation Suite^85^ with a human codon table obtained from the Codon Usage Database. Results were downloaded, sequences codon optimized using Geneious Prime (Biomatters) and stored. For germline reversion the most probable V gene derived by igblastp (igblast 1.16.0 package) was used as a template and V gene mutations from FWR1 to FWR3 were reverted, while the original antibody CDR3 and FWR4 regions were retained.

#### Antibody sequence analysis

For mutation analyses, mutations were counted based on the most probable V gene as derived by igblastp (igblast 1.16.0 package) and mutations were counted from FWR1 to FWR3 excluding any mutations in the original antibody CDR3 and FWR4 regions. Sequence alignments between wild-type and germline versions were generated using the PairwiseAligner object implemented in biopython (1.77) using the BLOSUM62 substitution matrix, open_gap_score of −10 and extend_gap_score of −0.5. All alignment mismatches were counted as mutations. Using the CDR and FWR region boundaries annotated by igblastn mutations were matched to their respective region. Antibody sequences were grouped into clonal clusters by the same VH gene and a minimal CDRH3 identity (defined by Levenshtein distance in relation to the length of the shorter CDRH3) of >=75%. CDRH3 hydrophobicity was calculated based on the Eisenberg-scale.^86^ Multiple sequence alignments were calculated with Clustal Omega (version 1.2.3) and logo plots generated using matplotlib (v3.3.4) implemented in a custom python script.

#### Cloning and production of antibody heavy and light chain

Heavy and light chain sequences of corresponding wild-type and germline antibodies were ordered as eBlocks gene fragments (IDT) including overhangs designed for subsequent cloning into expression vectors (IgG1, Igλ, Igκ).^87^ Sequence- and Ligation-Independent Cloning (SLIC) was performed using T4 DNA polymerase (New England Biolabs) and chemical competent *Escherichia coli* DH5α as previously described with a minor modification.^36,88–90^ In contrast to the published protocol a murine leader sequence encoded in the vector was used for secretion. Colony PCR and subsequent Sanger sequencing was used to verify positive colonies. Bacterial cultures of positive colonies were inoculated in LB-Medium, and plasmids isolated using the NucleoBond Xtra Midi kit (Macherey-Nagel). Monoclonal antibodies were produced in 50 ml of 0.8×10^6^ HEK293-6E cells by co-transfection of heavy chain (IgG1) and light chain (Igλ, Igκ) antibody expression plasmids with polyethylenimine (PEI; Sigma-Aldrich). Cells were maintained at 37°C and 5% CO_2_ in FreeStyle 293 Expression Medium (Thermo-Fisher) and 0.2% penicillin/streptomycin (Thermo-Fisher) under constant shaking at 120 rpm. Supernatants were harvested by centrifugation 7 days after transfection and incubated with Protein G-coupled beads overnight at 4°C (GE Life Sciences). Beads were loaded onto chromatography columns (Bio-Rad), washed with DPBS (Thermo-Fisher) and antibodies eluted using 0.1 M glycine (pH = 3) into 1 M Tris (pH = 8). 30K Amicon spin membranes (Merck Milipore) were used to exchange buffer to PBS for the 50ml transfections. Final antibody concentration was determined using UV spectrophotometry (Nanodrop, Thermo-Fisher).

#### Cloning and production of SARS-CoV-2 spike protein

The following coronavirus regions were amplified from synthetic gene plasmids and cloned into modified Sleeping Beauty transposon expression vectors:^91^ Wu01 spike (MN908947; AA: 1-1207; RRAR to GGGG; K986P; V987P; C-terminal T4 foldon – Twin-Strep-tag; 139 kDa); SARS-CoV-2 HexaPro BA.1 spike (MN908947; AA: 16-1208, furin site: RRAR to GSAS, A76V, delta69-70, T95I, G142D, delta143-145, N211I, delta212, 215EPEins, G339D, S371L, S373P, S375F, K417N, N440K, G446S, S477N, T478K, E484A, Q493R, G496S, Q498R, N501Y, Y505H, T547K, D614G, H655Y, N679K, P681H, N764K, D796Y, N856K, Q954H, N969K, L981F, including the stabilizing mutations: F817P, A892P, A899P, A942P, K986P, V987P, N-terminal BM40 signal peptide, C-terminal T4 foldon followed by a Twin strep tag, 139 kDa); SARS-CoV-2 Delta S (MN908947: AA:1-1207; RRAR to GGSG; T19R, G142D, R156G, delta157-158, L450R, T476K, D612G, P679R, D948N, K986P, V987P, N-terminal BM40 signal peptide, C-terminal T4 foldon followed by a Twin strep tag, 139 kDa); OC43 spike (AAX84792; AA: 1-1300; RRSRR to GSAS; A1078P; L1079P; C-terminal T4 foldon – Twin-Strep-tag; 150 kDa); 229E S1 (BAL45639; AA: 22-539, N-terminal BM-40 signal peptide – Twin-Strep-tag; 61 kDa) HKU1 S1 (YP_173238; AA: 14-612, N-terminal BM-40 signal peptide – Twin-Strep-tag; 72 kDa) and NL63 S1 (AKT07952; AA: 16-619, N-terminal BM-40 signal peptide – Twin-Strep-tag; 71 kDa). For recombinant protein production, stable HEK293 EBNA cell lines were generated using the sleeping beauty transposon system.^91^ Expression constructs were co-transfected with a transposase plasmid (10:1) into the HEK293 EBNA cells using FuGENE® HD transfection reagent (Promega GmbH, Madison, USA) in DMEM/F12 supplemented with 6% FBS. After high puromycin selection (3 µg/ml; Sigma-Aldrich), cells were expanded in triple flasks and protein production induced with doxycycline (0.5 µg/ml, Sigma-Aldrich). Supernatants of confluent cells were harvested every 3 days, filtered and recombinant proteins purified via Strep-Tactin®XT (IBA Lifescience, Göttingen, Germany) resin. Proteins were eluted with biotin containing buffer (IBA Lifescience, Göttingen, Germany), dialyzed against TBS and stored at 4°C or −80°C.

#### ELISA analysis for binding activity to SARS-CoV-2 spike protein

High affinity ELISA plates (Corning 3369) were coated at 4°C overnight with 2 µg/ml of SARS-CoV-2 spike protein in PBS (Thermo-Fisher). Plates were blocked using 5% milk in PBS for 60 min at room temperature and then washed using PBST (PBS supplemented with 0.2% Tween (Carl-Roth)). Primary antibodies were serially diluted in PBS, transferred onto the ELISA plates, and incubated for 90 min at room temperature followed by another washing step with PBST. The secondary antibody, anti-human IgG-HRP (Southern Biotech 2040-05), was diluted 1:2500 in PBS, transferred onto the ELISA plates and incubated for 60 min followed by the last washing step using PBST. ELISAs were developed using ABTS solution (Thermo Fisher) and absorbance was measured at 415 nm and 695 nm by a plate reader (Tecan). Background signal was subtracted from measured OD values and positive binding was defined by an OD ≥ 0.25. The area under the curve (AUC) was calculated using Prism 9 (total peak area above baseline). AUC values were normalized to the AUC of a control antibody (HbnC3t1p2_B10)^72^ running on every plate. For human coronavirus ELISA AUC was normalized on the AUC of Streptactin-HRP (IBA Lifescience) running on every plate. Binding levels of monoclonal antibodies (EC_50_) were calculated using a non-linear fit model (agonist vs response – variable slope (four parameters)).

#### SARS-CoV-2 pseudo-typed lentivirus cloning and production

All SARS-CoV-2 spike constructs were expressed using codon-optimized lentiviral expression plasmids. SARS-CoV-2 Wu01 spike (EPI_ISL_40671), Alpha, Beta, Delta and Omicron sublineages BA.4/5 were cloned in a pCDNA^TM^3.1/V5-HisTOPO vector and pseudo-typed lentivirus was produced as described previously.^92^ In brief, individual plasmids encoding HIV-1 Tat, HIV-1 Gag/Pol, HIV-1 Rev, Firefly luciferase followed by an IRES-ZsGreen, and the corresponding SARS-CoV-2 spike construct were co-transfected in HEK293-T cells using FuGENE 6 Transfection Reagent (Promega) in Dulbecco’s Modified Eagle Medium (DMEM, Thermo Fisher). Culture supernatant was harvested at 48 h and 72 h post transfection and stored at −80°C.

#### Pseudo-typed virus neutralization assays

The pseudo-typed virus neutralization assay was performed as described previously^92^. In summary, titrated pseudo-typed virus supernatants were co-incubated with a serial dilution of monoclonal antibodies at 37°C and 5 % CO_2_ for 60 min prior to the addition of HEK293-T cells stably expressing ACE2. Following a 48 h incubation period at 37°C and 5% CO_2_, relative light units (RLU) were determined using a microplate reader (Berthold) after the addition of luciferin/lysis buffer (10 mM MgCl2, 0.3 mM ATP, 0.5 mM coenzyme A, 17 mM IGEPAL CA-630 (all Sigma-Aldrich) and 1 mM D-Luciferin (GoldBio) in Tris-HCL). Average background RLU from uninfected control wells were subtracted and the 50% inhibitory concentrations (IC_50_s) were determined as the monoclonal antibody concentration that resulted in a 50% signal reduction compared to untreated infected controls, using a non-linear fit model (agonist vs normalized response curve with variable slope, GraphPad Prism).

#### Protein expression and purification for structural analysis of HbnC3t1p1_C6

The plasmid encoding the His-tagged SARS-CoV-2 receptor binding domain (RBD, AA 319-541 NM908947.3) was kindly provided by Florian Krammer.^93^ HbnC3t1p1_C6 IgG was expressed in HEK293F suspension cells (Invitrogen) by co-transfecting the heavy and light chains encoding vectors. Transfections were carried out using PEI-MAX (Polysciences). Media was collected at 6 days post-transfection, and IgG was captured using protein-A affinity chromatography (GE Healthcare). For obtaining the Fab, IgG was digested using Papain (Sigma-Aldrich) with enzyme to protein ratio being 1:1000. Digestion proceeded overnight in 16°C in a buffer containing 20 mM Cysteine-HCl (Sigma-Aldrich) and 10 mM EDTA titrated to pH = 7.0. Fabs were separated from Fc fragments by collecting the flow-through fraction from a protein-A column, followed by size exclusion chromatography (SEC) on Superdex 200 10/300 column (GE Healthcare). His-tagged SARS-CoV2 RBD was expressed in HEK293F cells as described above. Media was collected and buffer exchanged to PBS using a tangential flow filtration system (Millipore). Protein was captured using a HiTrap IMAC FF Ni^2+^ (GE Healthcare) affinity column followed by SEC purification with a Superdex 200 10/300 column. HbnC3t1p1_C6 Fab and SARS2 RBD complexes were created by mixing them at 1:1.2 molar ratio respectively.

#### Structural analysis of HbnC3t1p1_C6 antibody

For protein complexes crystallization we used a mosquito crystallization robot (TTP Labtech) to set vapor diffusion sitting drops with 96-well iQ plates (TTP Labtech), for each well, we tested three ratios of protein (80, 120 and 160 nl) to reservoir (120 nl). PEGrx-HT screen (Hampton Research) and ProPlex screen-HT (Molecular-dimensions) were used to identify initial hits which were obtained for apo-Fab HbnC3t1p1_C6 and Fab HbnC3t1p1_C6 bound to RBD respectively. Crystal hits were obtained for Apo-Fab HbnC3t1p1_C6 using protein concentration of 10.7 mg/ml and a protein reservoir ratio of 1:1. Optimized conditions contained 12% isopropanol, 0.08 M sodium citrate tribasic dihydrate and 22% polyethylene glycol 3350. Protein of Fab HbnC3t1p1_C6 in complex with RBD gave crystal hits in 0.1 M Tris pH = 8.5, 20% polyethylene glycol 6000 using 14 mg/ml protein in a 1.3:1 protein to reservoir ratio. 25% and 33% of ethylene glycol reservoir solution was used as a cryo-protectant for apo-Fab HbnC3t1p1_C6 and Fab HbnC3t1p1_C6 bound to RBD respectively. All crystals were grown in 20°C incubation.

X-ray data collection, structure determination and refinement. X-ray diffraction data for Fab HbnC3t1p1_C6 in complex with SARS-CoV-2 RBD were collected using Rigoku R-axis IV++ home source at 3 Å resolution from a crystal belonging to an orthorhombic space group. Data was further indexed, integrated, and scaled using HKL2000.^94^ We used Phaser^95^ to obtain a molecular replacement solution with a structure of a Fab (PDB 6XUK), and of the SARS-CoV-2 RBD (PDB 6M17) search models. The model was manually traced into electron density maps using Coot^96^ and refined using Phenix Refine.^97^ For generating the overlay in Figure 2 we used the following structures: 7B0B, 7E3K, 7E3L, 7BEN, 7P40, 7EZV, 7LRS.

## QUANTIFICATION AND STATISTICAL ANALYSIS

Statistical analyses were performed using Prism 9.0 (GraphPad), Microsoft Excel for Mac (v14.7.3) and Python (v3.6.8). In Supplementary Figure 1 a simple linear regression was used to determine the correlation between EC_50_/IC_50_ fold change and number of amino acid mutations. For the phylogenetic analyses in Figure 3 and Supplementary Figure 3 the amino acid sequences of the respective coronavirus spike proteins were aligned using Geneious Prime (Biomatters) and the tree was build using PhyML implemented in Geneious Prime (Biomatters). A p-value below 0.05 was considered significant. Additional details are provided in the figure legends.

