## Supplementary Information Korenkov et al. for "Dissecting the impact of somatic hypermutation on SARS-CoV-2 neutralization and viral escape"

### Supplementary Figures


| | |
|---|---|
| Supplementary Figure 1 | Sequence features of the 92 selected antibodies |
| Supplementary Figure 2 | Correlation of change in binding or neutralization after germline reversion with the number of reverted mutations |
| Supplementary Figure 3 | Cross-reactivity of wild-type and germline-reverted antibodies with endemic human coronavirus spike proteins |
| Supplementary Figure 4 | Correlation of change in neutralization after germline reversion with CDRH3 length and hydrophobicity |
| Supplementary Figure 5 | The effect of SHM on binding and neutralization potency grouped by VH and VL genes |
| Supplementary Figure 6 | Crystal structure of HbnC3t1p1_C6 and role of the disulfide bridge in the CDRH3 |
| Supplementary Figure 7 | The impact of SHM on binding of VH1-58 antibodies to Wu01, Delta, and Omicron BA.1 spike protein |
| Supplementary Figure 8 | The impact of grouped and individual mutation reversions on Omicron neutralizing VH1-58 antibodies |


### Supplementary Tables


| | |
|---|---|
| Supplementary Table 1 | Selected antibodies and sequence features |
| Supplementary Table 2 | Fab HbnC3t1p1_C6 - PDB entry 7B0B |
| Supplementary Table 3 | Initial and additional IGHV1-58/IGKV3-20 antibodies |

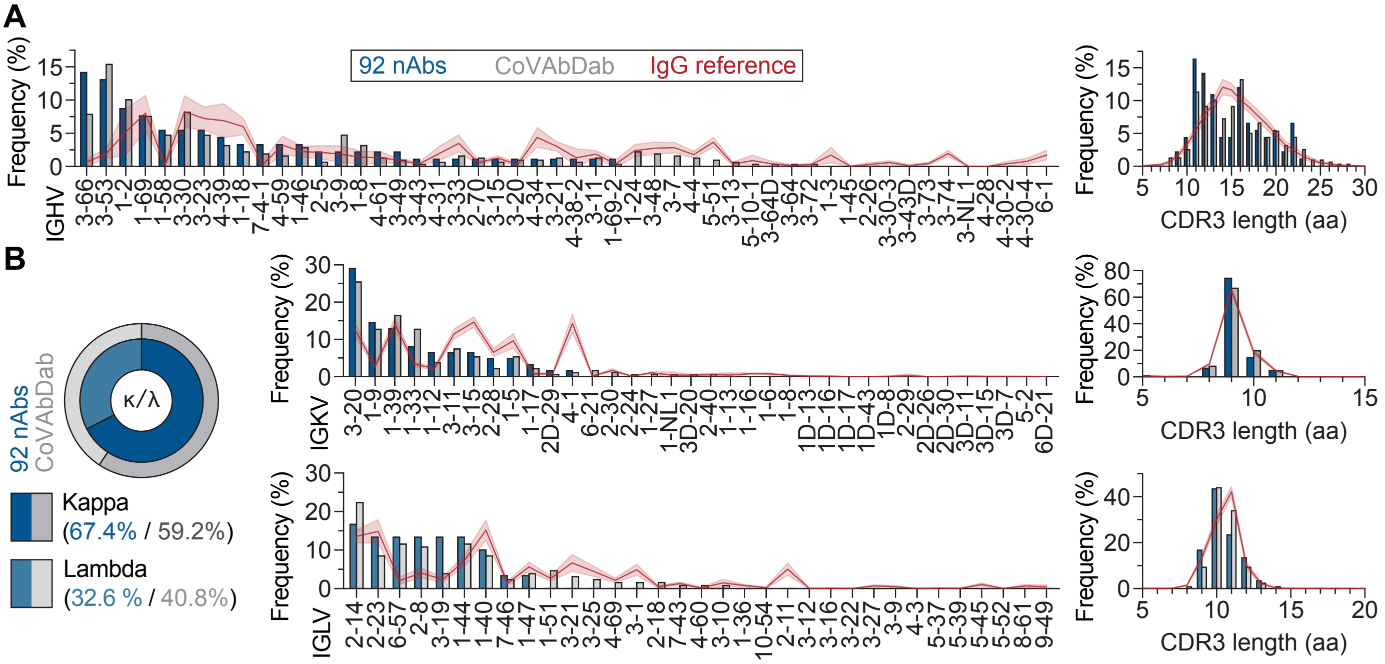


**Supplementary Figure 1: Sequence features of the 92 selected antibodies.** (**A**) Heavy chain V gene segment usage and complementarity determining region (CDR) 3 length distributions from the selected 92 antibodies in comparison to the complete 319 human SARS-CoV-2 neutralizing antibodies from the CoV-AbDab and IgG reference repertoires from n=57 healthy individuals. (**B**) Kappa and lambda fraction of light chains from the 92 selected antibodies and the CoV-AbDab antibodies, as well as V gene segment usage and CDR3 length distributions from the selected antibodies, the CoV-AbDab antibodies, and IgG reference repertoires.

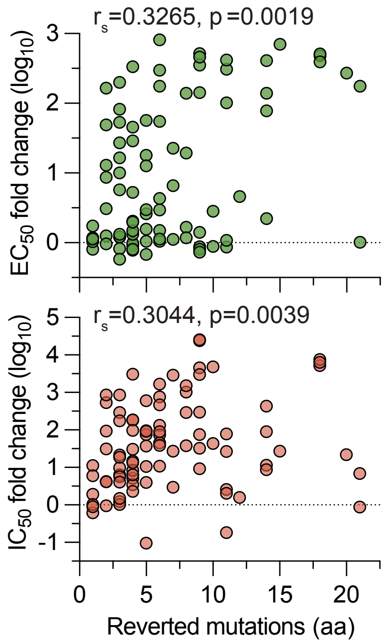


**Supplementary Figure 2: Correlation of change in binding or neutralization after germline reversion with the number of reverted mutations.** $Log_{10}$ fold changes in $EC_{50}$ (binding) or $IC_{50}$ (neutralization) after germline-reversion are plotted against the total number of reverted heavy and light chain mutations. Spearman correlation coefficients $r_s$ as well as corresponding p values are depicted in the plots. aa: amino acid.

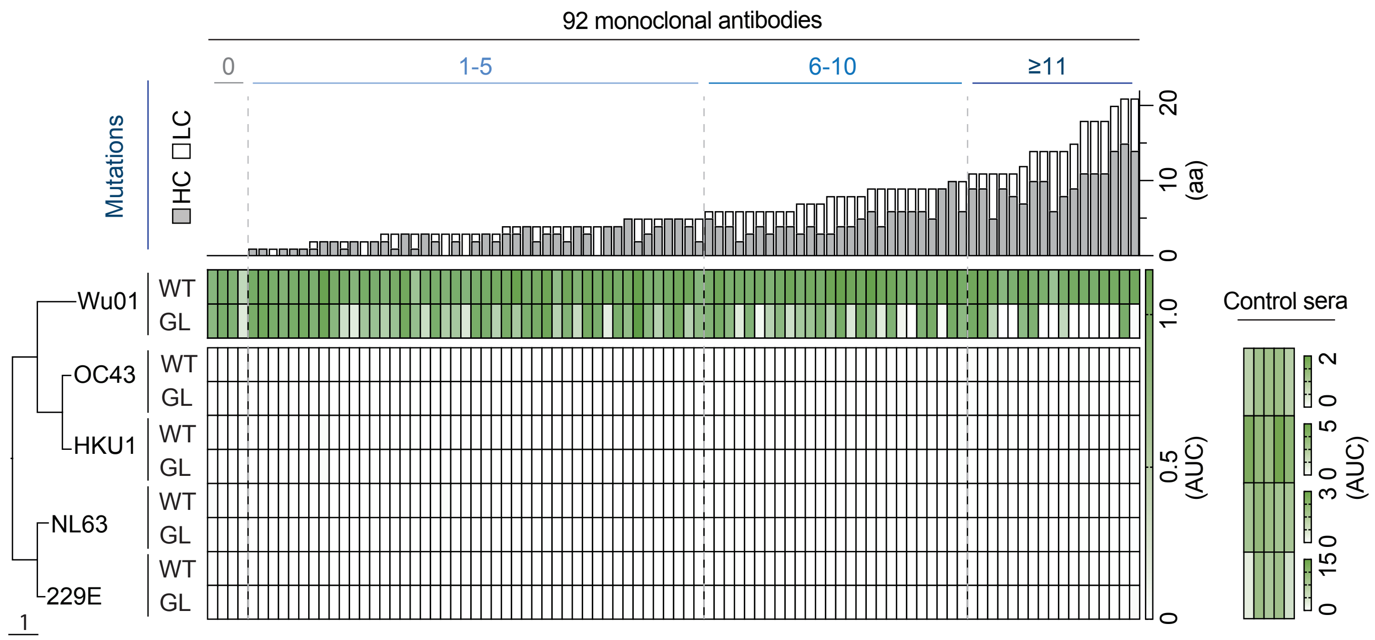


**Supplementary Figure 3: Cross-reactivity of wild-type and germline-reverted antibodies with endemic human coronavirus spike proteins.** Area under the curve (AUC) heatmap of ELISAs against Wu01, OC43, HKU1, 229E, and NL63 spike proteins. Human coronaviruses are ordered by their phylogenetic distance on amino acid (aa) level. The tree scale represents one amino acid substitution per site. Five control sera of healthy individuals were taken as positive controls for human coronavirus spike proteins. The upper part including the mutation count and Wu01 AUC data for individual wild-type and germline-reverted antibodies was taken from Figure 1D for comparison.

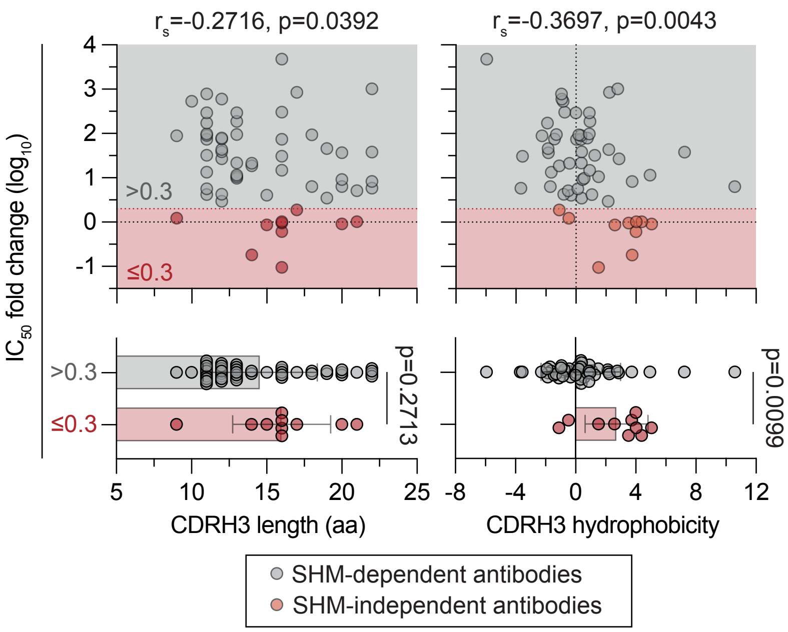


**Supplementary Figure 4: Correlation of change in neutralization after germline reversion with CDRH3 length and hydrophobicity.** $Log_{10}$ fold change $IC_{50}$ (neutralization) after germline-reversion is plotted against the CDRH3 length (aa: amino acids) and hydrophobicity (based on the Eisenberg scale) for antibodies that contained at least one mutation and did not lose neutralizing activity completely after germline reversion (n=58). Boundary for antibodies to count as unaffected by reversion was set to a $log_{10}$ fold change ≤0.3. Spearman correlation coefficients $r_s$ and corresponding p values as well as p values from two-tailed, unpaired t tests are depicted in the plots. CDR: complementarity determining region.

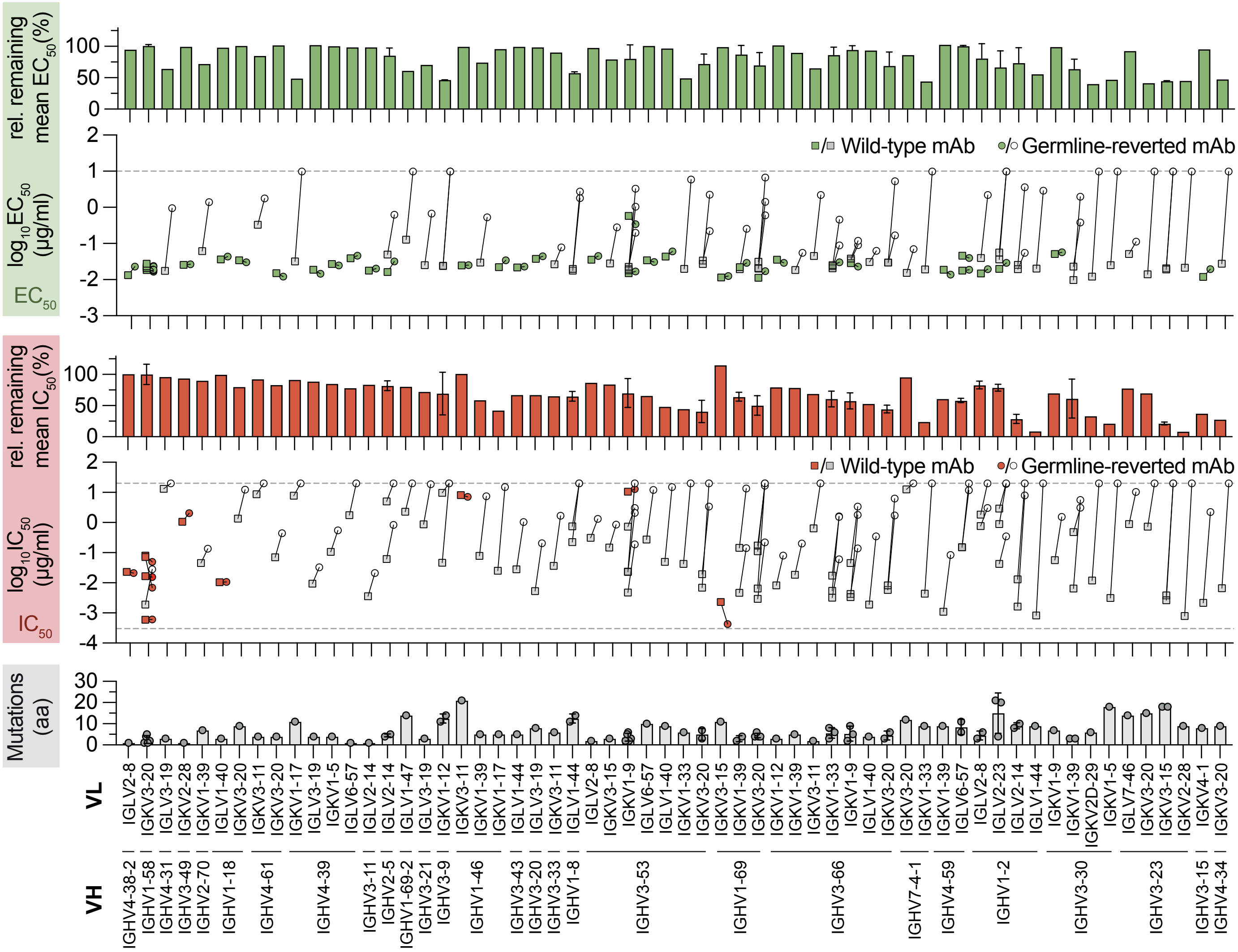


**Supplementary Figure 5: The effect of SHM on binding and neutralization potency grouped by VH and VL genes.** Monoclonal antibodies were grouped by VH and VL genes and sorted by the mean $log_{10}$ fold $IC_{50}$ change in each group as in Figure 2. Upper panel: Bars depict the mean remaining $EC_{50}$ after germline reversion normalized to the maximum detectable difference, i.e., lower limit of quantification (LLOQ, 0.0001 µg/ml) to upper limit of quantification (ULOQ, 10 µg/ml). Scatter plots show $log_{10}$ $EC_{50}$ values of individual antibodies before (squares) and after (circles) germline-reversion with corresponding pairs connected by a line. ULOQ is depicted as a dashed line. Antibodies that remained unaffected by reversion ($log_{10}$ fold $EC_{50}$ change ≤ 0.3) are highlighted in green. Middle panel: Bars depict the mean remaining IC50 after germline reversion normalized to the maximum detectable difference, i.e., LLOQ (0.0003 µg/ml) to ULOQ (20 µg/ml). Scatter plots show $log_{10}IC_{50}$ values of individual antibodies before (squares) and after (circles) germline-reversion with corresponding pairs connected by a line. ULOQ/LLOQ are depicted as dashed lines. Antibodies that remained unaffected by reversion ($log_{10}$ fold $IC_{50}$ change ≤ 0.3) are highlighted in red. Lower panel: Individual (dots) and mean (bars) number of mutations in the VH/VL group. Error bars depict standard deviation, where applicable.

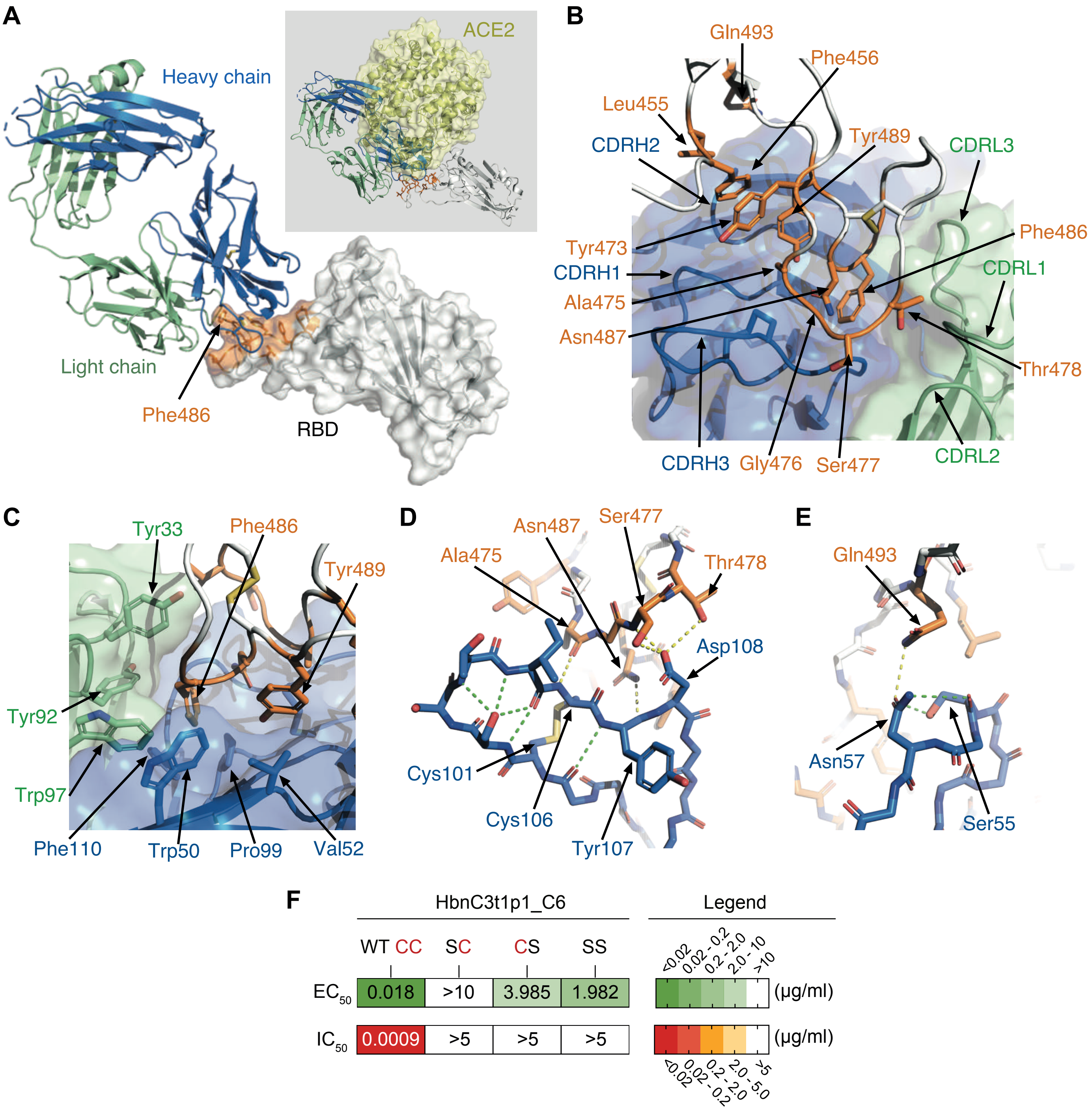


**Supplementary Figure 6: Crystal structure of HbnC3t1p1_C6 and role of the disulfide bridge in the CDRH3.** (**A**) The overall structure of the HbnC3t1p1_C6 Fab (blue and green for the heavy and light chains, respectively) in complex with the SARS-CoV-2 RBD (grey, semitransparent surface representation). The epitope of HbnC3t1p1_C6 is highlighted in orange, and Phe486, which is a central feature of the complex, is marked. The greyed inset shows the HbnC3t1p1_C6/RBD complex superimposed with the ACE2/RBD structure (PDB ID: 6M17). HbnC3t1p1_C6 and ACE2 substantially overlap with each other, indicating that neutralization is achieved by blocking the interaction with ACE2. (**B**) An overview of the HbnC3t1p1_C6 epitope. The residues of SARS-CoV-2 RBD that make the epitope of HbnC3t1p1_C6 are labeled and shown as orange sticks. The heavy and light chains of HbnC3t1p1_C6 are shown using a semitransparent surface in blue and green, respectively. The CDRs are labeled. (**C**) HbnC3t1p1_C6 makes a hydrophobic pocket that interacts with Phe486 of the RBD at the interface between its heavy and light chains. Tyr33 from CDRL1, Tyr92, and Trp97 from CDRL3 of the light chain make part of the hydrophobic pocket and are indicated. At the heavy chain, Pro99 at the base and Phe110 at the end of CDRH3 make the bottom of the pocket. The side of the pocket is made of Trp50, a framework residue of the heavy chain. Val52 of CDRH2 makes hydrophobic interaction with Tyr489 of the RBD. (**D**) CDRH3 makes polar interactions with RBD. The main chain of CDRH3 is shown as sticks. Cys106 and Cys101 make a disulfide bridge that, together with a series of hydrogen bonds (green dashed lines) that involve mainchain atoms, stabilize the conformation of CDRH3. Asp108, through its sidechain, and Cys106 and Tyr107, through their mainchain, make hydrogen bonds (yellow dashed lines) with Thr478, Ser477, Asn487, and Ala475 of the RBD. (**E**) CDRH2 makes polar interaction with RBD. Asn57 is stabilized by hydrogen bonds (green dashed lines) to the nearby Ser55 and further forms a hydrogen bond (yellow dashed line) with Gln493 of RBD. (**F**) Binding and neutralization of HbnC3t1p1_C6 with single (SC, CS) and double (SS) cysteine substitutions to serine.

| $V_H$1-58 mAb | Wu01 WT | Wu01 GL | Delta WT | Delta GL | BA.1 WT | BA.1 GL |
|---|---|---|---|---|---|---|
| C043 | 0.007 | 0.012 | 0.006 | 0.010 | 0.007 | >10 |
| CQTS004 | 0.016 | 0.013 | 0.013 | 0.011 | 0.039 | >10 |
| C827 | 0.019 | 0.026 | 0.017 | 0.022 | 0.020 | 0.074 |
| R200-1B9 | 0.035 | 0.041 | 0.017 | 0.014 | 0.005 | 0.046 |
| R259-1B9 | 0.021 | 0.028 | 0.013 | 0.035 | 0.026 | >10 |
| Ab_58G6 | 0.017 | 0.014 | 0.014 | 0.013 | 0.057 | 0.920 |
| Turner-07-2C08 | 0.143 | 0.112 | 0.135 | 0.102 | 0.255 | >10 |
| C598 | 0.020 | 0.011 | 0.018 | 0.010 | 0.024 | 1.380 |
| COV2-2072 | 0.015 | 0.068 | 0.013 | 0.058 | 0.307 | 2.327 |
| HbnC3t1p1_C6 | 0.021 | 0.023 | 0.010 | 0.008 | 0.507 | 0.836 |
| COV2-3025 | 0.020 | 0.012 | 0.017 | 0.011 | >10 | 1.661 |
| I14 | 0.020 | 0.046 | 0.018 | 0.043 | >10 | >10 |
| COV2-2196 | 0.016 | 0.016 | 0.012 | 0.008 | 1.599 | >10 |
| P008_086 | 0.015 | 0.032 | 0.015 | 0.034 | >10 | >10 |
| C597 | 0.072 | 0.019 | 0.050 | 0.017 | >10 | >10 |
| Wang-C387 | 0.010 | 0.005 | 0.012 | 0.006 | >10 | >10 |
| COV2-2381 | 0.027 | 0.025 | 0.024 | 0.020 | >10 | >10 |
| HbnC3t1p2_C6 | 0.023 | 0.022 | 0.017 | 0.006 | >10 | >10 |
| C125 | 0.012 | 0.038 | 0.011 | 0.033 | >10 | >10 |
| P008_081 | 0.047 | 0.019 | 0.049 | 0.019 | >10 | >10 |
| MnC5t2p1_G1 | 0.027 | 0.019 | 0.012 | 0.010 | 2.538 | >10 |
| CV07-287 | 0.043 | 0.021 | 0.038 | 0.045 | >10 | >10 |

$EC_{50}$ (µg/ml): <0.02 | 0.02 - 0.2 | 0.2 - 2.0 | 2.0 - 10 | >10

**Supplementary Figure 7: The impact of SHM on binding of VH1-58 antibodies to Wu01, Delta, and Omicron BA.1 spike protein.** $EC_{50}$ values were determined by ELISA against Wu01, Delta, and Omicron BA.1 trimeric spike protein for original wild-type (WT) and germline-reverted (GL) antibodies. Values represent biological duplicates. Red labels depict Omicron BA.1 and BA.2 neutralizing antibodies.

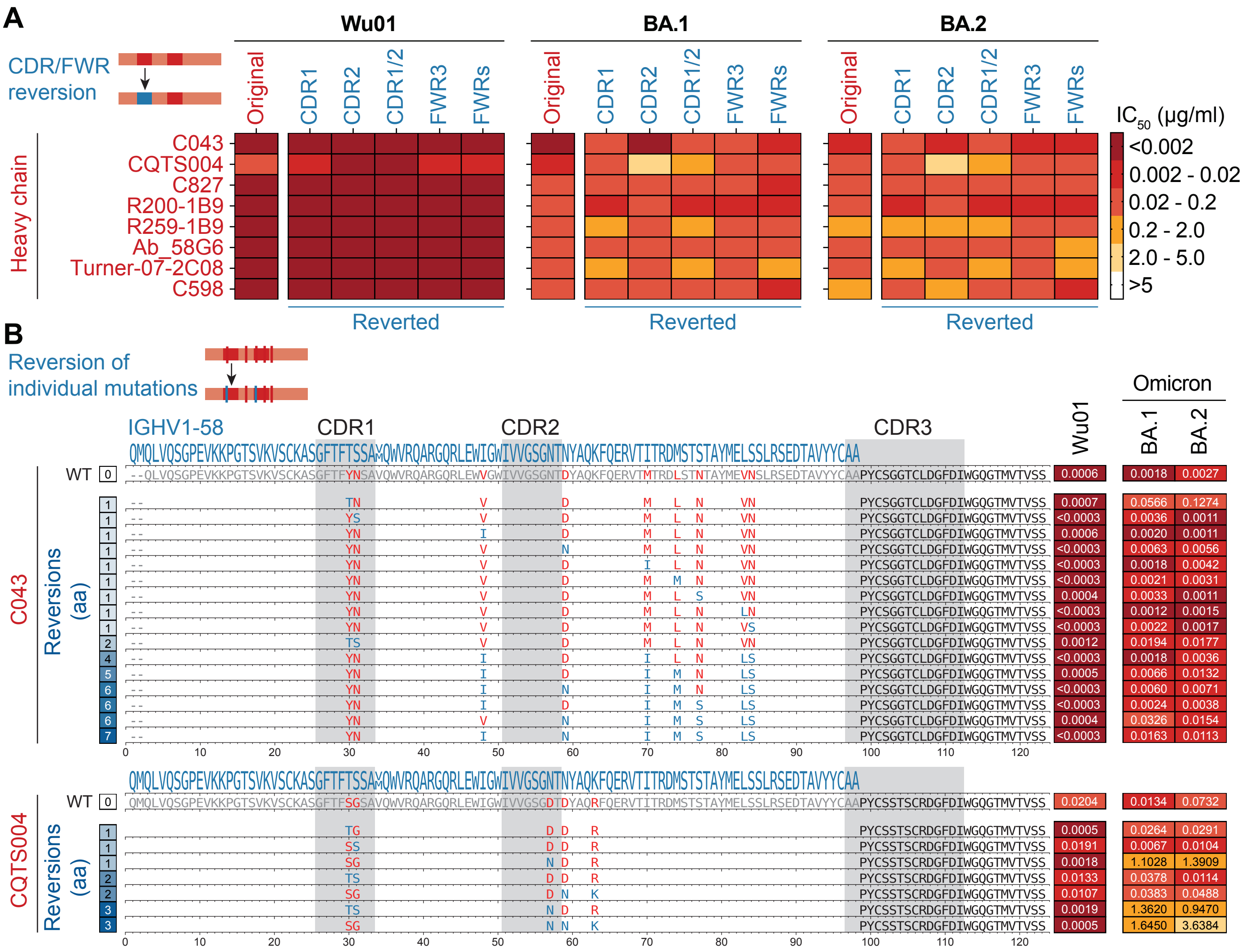

**Supplementary Figure 8: The impact of grouped and individual mutation reversions on Omicron neutralizing VH1-58 antibodies.** (**A**) Heavy chain CDRs or FWRs were reverted to germline in eight Omicron neutralizing antibodies and partially reverted antibodies (paired with original unreverted light chains) were tested for neutralization against Wu01, BA.1, and BA.2. Heatmaps depict mean $IC_{50}$ values for partially reverted antibodies from two independent experiments. (**B**) Additional individual or combinations of heavy chain mutations were reverted for CQTS004 and C043 as illustrated by blue amino acids in the sequence alignment and $IC_{50}$ values were determined. $IC_{50}$ values represent means of two independent experiments. Reversion variants in (B) are sorted by the number of reverted amino acids. Original wild-type (WT) neutralization values were taken from Figure 3A. CDR: complementarity determining region; FWR: framework region; aa: amino acid.

**Supplementary Table 1: Selected antibodies and sequence features**

| # | Name | Total mutations (aa) | VH gene | VH mutations (aa) | CDRH3 length (aa) | VL gene | VL mutations (aa) | CDRL3 length (aa) | Origin | Reference |
|---|---|---|---|---|---|---|---|---|---|---|
| 1 | CnC2t1p1_B4 | 0 | IGHV1-18 | 0 | 12 | IGLV2-23 | 0 | 10 | SARS-CoV-2 convalescent - memory B cell (IgG) | Kreer et al., Cell, 2020 |
| 2 | CnC2t1p1_D6 | 0 | IGHV3-49 | 0 | 17 | IGKV2-28 | 0 | 9 | SARS-CoV-2 convalescent - memory B cell (IgG) | Kreer et al., Cell, 2020 |
| 3 | MnC2t1p1_C5 | 0 | IGHV3-66 | 0 | 13 | IGKV1-12 | 0 | 9 | SARS-CoV-2 convalescent - memory B cell (IgG) | Kreer et al., Cell, 2020 |
| 4 | BD23 | 0 | IGHV7-4-1 | 0 | 19 | IGKV1-5 | 0 | 9 | SARS-CoV-2 convalescent - memory B cell (IgG) | Cao et al., Cell, 2020 |
| 5 | C125 | 1 | IGHV1-58 | 1 | 16 | IGKV3-20 | 0 | 9 | SARS-CoV-2 convalescent - memory B cell (IgG) | Robbiani et al., Nature, 2020 |
| 6 | CnC2t1p1_E12 | 1 | IGHV3-49 | 1 | 17 | IGKV2-28 | 0 | 9 | SARS-CoV-2 convalescent - memory B cell (IgG) | Kreer et al., Cell, 2020 |
| 7 | HbnC3t1p1_C6 | 1 | IGHV1-58 | 0 | 16 | IGKV3-20 | 1 | 9 | SARS-CoV-2 convalescent - memory B cell (IgG) | Kreer et al., Cell, 2020 |
| 8 | COV2-2677 | 1 | IGHV4-39 | 1 | 12 | IGLV6-57 | 0 | 10 | SARS-CoV-2 convalescent - class switched memory B cell | Zost et al., Nature Medicine, 2020 |
| 9 | CV07-270 | 1 | IGHV3-11 | 1 | 22 | IGLV2-14 | 0 | 10 | SARS-CoV-2 convalescent - memory B cell (IgG) | Kreye et al., Cell, 2020 |
| 10 | P2B-2F6 | 1 | IGHV4-38-2 | 1 | 20 | IGLV2-8 | 0 | 10 | SARS-CoV-2 convalescent - memory B cell (IgG) | Ju et al., Nature, 2020 |
| 11 | C105 | 2 | IGHV3-53 | 1 | 12 | IGLV2-8 | 1 | 11 | SARS-CoV-2 convalescent - memory B cell (IgG) | Robbiani et al., Nature, 2020 |
| 12 | COV2-2196 | 2 | IGHV1-58 | 2 | 16 | IGKV3-20 | 0 | 10 | SARS-CoV-2 convalescent - class switched memory B cell | Zost et al., Nature Medicine, 2020 |
| 13 | CV38-139 | 2 | IGHV3-66 | 2 | 10 | IGKV1-9 | 0 | 10 | SARS-CoV-2 convalescent - memory B cell (IgG) | Kreye et al., Cell, 2020 |
| 14 | DH1197 | 2 | IGHV3-66 | 1 | 14 | IGKV3-11 | 1 | 10 | SARS-CoV-2 convalescent - class switched memory B cell | Li et al., Cell, 2021 |
| 15 | C123 | 2 | IGHV3-53 | 2 | 10 | IGKV1-9 | 0 | 9 | SARS-CoV-2 convalescent - memory B cell (IgG) | Robbiani et al., Nature, 2020 |
| 16 | CV-X2-106 | 2 | IGHV1-69 | 0 | 18 | IGKV1-39 | 2 | 9 | SARS-CoV-2 convalescent - memory B cell (IgG) | Kreye et al., Cell, 2020 |
| 17 | C210 | 2 | IGHV3-53 | 2 | 11 | IGKV1-9 | 0 | 10 | SARS-CoV-2 convalescent - memory B cell (IgG) | Robbiani et al., Nature, 2020 |
| 18 | COVA2-29 | 3 | IGHV3-30 | 2 | 18 | IGKV1-39 | 1 | 9 | SARS-CoV-2 convalescent - single B cell | Brouwer et al., Science, 2020 |
| 19 | HbnC3t1p1_G4 | 3 | IGHV3-66 | 1 | 11 | IGKV3-20 | 2 | 9 | SARS-CoV-2 convalescent - memory B cell (IgG) | Kreer et al., Cell, 2020 |
| 20 | MnC2t1p1_A3 | 3 | IGHV3-66 | 3 | 13 | IGKV1-12 | 0 | 9 | SARS-CoV-2 convalescent - memory B cell (IgG) | Kreer et al., Cell, 2020 |
| 21 | B38 | 3 | IGHV3-53 | 1 | 9 | IGKV1-9 | 2 | 10 | SARS-CoV-2 convalescent - memory B cell | Wu et al., Science, 2020 |
| 22 | C102 | 3 | IGHV3-53 | 3 | 11 | IGKV3-20 | 0 | 9 | SARS-CoV-2 convalescent - memory B cell (IgG) | Robbiani et al., Nature, 2020 |
| 23 | S2X35 | 3 | IGHV1-18 | 2 | 21 | IGLV1-40 | 1 | 13 | SARS-CoV-2 convalescent - memory B cell (IgG) | Piccoli et al., Cell, 2020 |
| 24 | Fab2-38 | 3 | IGHV3-21 | 0 | 14 | IGLV3-19 | 3 | 9 | SARS-CoV-2 convalescent - memory B cell | Liu et al., Nature, 2020 |
| 25 | C155 | 3 | IGHV3-53 | 2 | 11 | IGKV3-15 | 1 | 9 | SARS-CoV-2 convalescent - memory B cell (IgG) | Robbiani et al., Nature, 2020 |
| 26 | DH1171 | 3 | IGHV4-31 | 0 | 15 | IGLV3-19 | 3 | 12 | SARS-CoV-2 convalescent - memory B cell or plasmablast | Li et al., Cell, 2021 |
| 27 | REGN10970 | 3 | IGHV3-66 | 2 | 14 | IGKV1-33 | 1 | 9 | SARS-CoV-2 convalescent - single B cell | Hansen et al., Science, 2020 |
| 28 | Fab2-4 | 3 | IGHV1-2 | 3 | 15 | IGLV2-8 | 0 | 10 | SARS-CoV-2 convalescent - memory B cell | Liu et al., Nature, 2020 |
| 29 | C002 | 3 | IGHV3-30 | 2 | 17 | IGKV1-39 | 1 | 9 | SARS-CoV-2 convalescent - memory B cell (IgG) | Robbiani et al., Nature, 2020 |
| 30 | C022 | 4 | IGHV4-39 | 3 | 21 | IGKV1-5 | 1 | 9 | SARS-CoV-2 convalescent - memory B cell (IgG) | Robbiani et al., Nature, 2020 |
| 31 | C165 | 4 | IGHV1-69 | 3 | 15 | IGKV3-20 | 1 | 9 | SARS-CoV-2 convalescent - memory B cell (IgG) | Robbiani et al., Nature, 2020 |
| 32 | HbnC3t1p2_C6 | 4 | IGHV1-58 | 4 | 16 | IGKV3-20 | 0 | 9 | SARS-CoV-2 convalescent - memory B cell (IgG) | Kreer et al., Cell, 2020 |
| 33 | COV2-2499 | 4 | IGHV4-39 | 2 | 19 | IGLV3-19 | 2 | 11 | SARS-CoV-2 convalescent - class switched memory B cell | Zost et al., Nature Medicine, 2020 |
| 34 | DH1042 | 4 | IGHV1-69 | 3 | 16 | IGKV1-39 | 1 | 9 | SARS-CoV-2 convalescent - memory B cell or plasmablast | Li et al., Cell, 2021 |
| 35 | DH1138 | 4 | IGHV4-61 | 3 | 12 | IGKV3-11 | 1 | 8 | SARS-CoV-2 convalescent - memory B cell or plasmablast | Li et al., Cell, 2021 |
| 36 | REGN10986 | 4 | IGHV3-66 | 2 | 13 | IGLV1-40 | 2 | 12 | SARS-CoV-2 convalescent - single B cell | Hansen et al., Science, 2020 |
| 37 | CC6.33 | 4 | IGHV1-69 | 4 | 11 | IGKV3-20 | 0 | 9 | SARS-CoV-2 convalescent - memory B cell (IgG) | Rogers et al., Science, 2020 |
| 38 | CV07-262 | 4 | IGHV1-2 | 3 | 22 | IGLV2-23 | 1 | 10 | SARS-CoV-2 convalescent - memory B cell (IgG) | Kreye et al., Cell, 2020 |
| 39 | Fab2-7 | 4 | IGHV2-5 | 0 | 11 | IGLV2-14 | 4 | 9 | SARS-CoV-2 convalescent - memory B cell | Liu et al., Nature, 2020 |
| 40 | DH1184 | 4 | IGHV1-69 | 4 | 18 | IGKV3-20 | 0 | 9 | SARS-CoV-2 convalescent - memory B cell or plasmablast | Li et al., Cell, 2021 |
| 41 | Fab2-36 | 4 | IGHV4-61 | 4 | 20 | IGKV3-20 | 0 | 9 | SARS-CoV-2 convalescent - memory B cell | Liu et al., Nature, 2020 |
| 42 | C140 | 5 | IGHV3-66 | 5 | 11 | IGKV1-9 | 0 | 9 | SARS-CoV-2 convalescent - memory B cell (IgG) | Robbiani et al., Nature, 2020 |
| 43 | MnC5t2p1_G1 | 5 | IGHV1-58 | 2 | 16 | IGKV3-20 | 3 | 9 | SARS-CoV-2 convalescent - memory B cell (IgG) | Kreer et al., Cell, 2020 |
| 44 | CB6 | 5 | IGHV3-66 | 3 | 13 | IGKV1-39 | 2 | 11 | SARS-CoV-2 convalescent - memory B cell (IgG) | Shi et al., Nature, 2020 |
| 45 | Fab4-20 | 5 | IGHV1-46 | 4 | 13 | IGKV1-39 | 1 | 10 | SARS-CoV-2 convalescent - memory B cell | Liu et al., Nature, 2020 |
| 46 | CC6.31 | 5 | IGHV1-46 | 5 | 12 | IGKV1-17 | 0 | 10 | SARS-CoV-2 convalescent - memory B cell (IgG) | Rogers et al., Science, 2020 |
| 47 | GW01 | 5 | IGHV3-43 | 5 | 20 | IGLV1-44 | 0 | 10 | SARS-CoV-2 convalescent - memory B cell (IgG) | CN111793129A, 2020; Wang et al., Cell Discovery, 2022 |
| 48 | REGN10971 | 5 | IGHV3-53 | 4 | 11 | IGKV1-9 | 1 | 10 | SARS-CoV-2 convalescent - single B cell | Hansen et al., Science, 2020 |
| 49 | COV2-2268 | 5 | IGHV2-5 | 2 | 11 | IGLV2-14 | 3 | 11 | SARS-CoV-2 convalescent - class switched memory B cell | Zost et al., Nature Medicine, 2020 |
| 50 | COV2-2841 | 6 | IGHV4-59 | 5 | 12 | IGLV6-57 | 1 | 9 | SARS-CoV-2 convalescent - class switched memory B cell | Zost et al., Nature Medicine, 2020 |
| 51 | REGN10977 | 6 | IGHV1-69 | 4 | 16 | IGKV3-20 | 2 | 9 | SARS-CoV-2 convalescent - single B cell | Hansen et al., Science, 2020 |
| 52 | HbnC2t1p2_D9 | 6 | IGHV3-33 | 4 | 19 | IGKV3-11 | 2 | 11 | SARS-CoV-2 convalescent - memory B cell (IgG) | Kreer et al., Cell, 2020 |
| 53 | HbnC3t1p2_B10 | 6 | IGHV3-66 | 2 | 11 | IGKV3-20 | 4 | 9 | SARS-CoV-2 convalescent - memory B cell (IgG) | Kreer et al., Cell, 2020 |
| 54 | BD-236 | 6 | IGHV3-53 | 3 | 12 | IGKV1-9 | 3 | 9 | SARS-CoV-2 convalescent - memory B cell (IgG) | Cao et al., Cell, 2020 |
| 55 | COV2-2955 | 6 | IGHV3-30 | 4 | 22 | IGKV2D-29 | 2 | 9 | SARS-CoV-2 convalescent - class switched memory B cell | Zost et al., Nature Medicine, 2020 |
| 56 | CC12.4 | 6 | IGHV1-2 | 3 | 19 | IGLV2-8 | 3 | 10 | SARS-CoV-2 convalescent - memory B cell (IgG) | Rogers et al., Science, 2020 |
| 57 | REGN10955 | 6 | IGHV3-66 | 4 | 9 | IGKV1-33 | 2 | 9 | SARS-CoV-2 convalescent - single B cell | Hansen et al., Science, 2020 |
| 58 | COV2-2752 | 6 | IGHV3-53 | 4 | 10 | IGKV1-33 | 2 | 9 | SARS-CoV-2 convalescent - class switched memory B cell | Zost et al., Nature Medicine, 2020 |
| 59 | COV2-2919 | 7 | IGHV2-70 | 3 | 12 | IGKV1-39 | 4 | 9 | SARS-CoV-2 convalescent - class switched memory B cell | Zost et al., Nature Medicine, 2020 |
| 60 | C101 | 7 | IGHV3-53 | 4 | 11 | IGKV3-20 | 3 | 9 | SARS-CoV-2 convalescent - memory B cell (IgG) | Robbiani et al., Nature, 2020 |
| 61 | Fab2-30 | 7 | IGHV3-30 | 3 | 12 | IGKV1-9 | 4 | 9 | SARS-CoV-2 convalescent - memory B cell | Liu et al., Nature, 2020 |
| 62 | REGN10954 | 8 | IGHV3-66 | 3 | 13 | IGKV1-33 | 5 | 9 | SARS-CoV-2 convalescent - single B cell | Hansen et al., Science, 2020 |
| 63 | COV2-2130 | 8 | IGHV3-15 | 4 | 22 | IGKV4-1 | 4 | 8 | SARS-CoV-2 convalescent - class switched memory B cell | Zost et al., Nature Medicine, 2020 |
| 64 | Fab2-15 | 8 | IGHV1-2 | 4 | 22 | IGLV2-14 | 4 | 10 | SARS-CoV-2 convalescent - memory B cell | Liu et al., Nature, 2020 |
| 65 | COV2-2678 | 8 | IGHV3-20 | 5 | 22 | IGLV3-19 | 3 | 11 | SARS-CoV-2 convalescent - class switched memory B cell | Zost et al., Nature Medicine, 2020 |
| 66 | COV2-2068 | 9 | IGHV3-53 | 6 | 16 | IGLV1-40 | 3 | 12 | SARS-CoV-2 convalescent - class switched memory B cell | Zost et al., Nature Medicine, 2020 |
| 67 | COV2-2050 | 9 | IGHV1-2 | 4 | 23 | IGLV1-44 | 5 | 11 | SARS-CoV-2 convalescent - class switched memory B cell | Zost et al., Nature Medicine, 2020 |
| 68 | S309 | 9 | IGHV1-18 | 6 | 20 | IGKV3-20 | 3 | 8 | SARS-CoV-1 convalescent - memory B cell | Pinto et al., Nature, 2020 |
| 69 | FnC1t2p1_D4 | 9 | IGHV7-4-1 | 6 | 11 | IGKV1-33 | 3 | 9 | SARS-CoV-2 convalescent - memory B cell | Kreer et al., Cell, 2020 |
| 70 | C104 | 9 | IGHV4-34 | 6 | 17 | IGKV3-20 | 3 | 9 | SARS-CoV-2 convalescent - memory B cell | Robbiani et al., Nature, 2020 |
| 71 | COV2-2952 | 9 | IGHV3-66 | 6 | 11 | IGKV1-9 | 3 | 9 | SARS-CoV-2 convalescent - class switched memory B cell | Zost et al., Nature Medicine, 2020 |
| 72 | REGN10964 | 9 | IGHV4-59 | 5 | 12 | IGKV1-39 | 4 | 9 | SARS-CoV-2 convalescent - single B cell | Hansen et al., Science, 2020 |
| 73 | BD-368-2 | 9 | IGHV3-23 | 9 | 18 | IGKV2-28 | 0 | 9 | SARS-CoV-2 convalescent - memory B cell | Cao et al., Cell, 2020 |
| 74 | COV2-2354 | 10 | IGHV3-53 | 10 | 12 | IGLV6-57 | 0 | 10 | SARS-CoV-2 convalescent - class switched memory B cell | Zost et al., Nature Medicine, 2020 |
| 75 | REGN10989 | 10 | IGHV1-2 | 6 | 16 | IGLV2-14 | 4 | 10 | SARS-CoV-2 convalescent - single B cell | Hansen et al., Science, 2020 |
| 76 | COV2-2531 | 11 | IGHV4-59 | 9 | 12 | IGLV6-57 | 2 | 9 | SARS-CoV-2 convalescent - class switched memory B cell | Zost et al., Nature Medicine, 2020 |
| 77 | COV2-2479 | 11 | IGHV1-69 | 9 | 14 | IGKV3-15 | 2 | 8 | SARS-CoV-2 convalescent - class switched memory B cell | Zost et al., Nature Medicine, 2020 |
| 78 | COV2-2539 | 11 | IGHV1-8 | 5 | 16 | IGLV1-44 | 6 | 11 | SARS-CoV-2 convalescent - class switched memory B cell | Zost et al., Nature Medicine, 2020 |
| 79 | MnC4t2p1_D10 | 11 | IGHV4-39 | 9 | 20 | IGKV1-17 | 2 | 9 | SARS-CoV-2 convalescent - memory B cell (IgG) | Kreer et al., Cell, 2020 |
| 80 | MnC4t2p1_E6 | 11 | IGHV3-9 | 8 | 17 | IGKV1-12 | 3 | 9 | SARS-CoV-2 convalescent - memory B cell (IgG) | Kreer et al., Cell, 2020 |
| 81 | MnC4t2p2_A4 | 12 | IGHV7-4-1 | 7 | 11 | IGKV3-20 | 5 | 9 | SARS-CoV-2 convalescent - memory B cell (IgG) | Kreer et al., Cell, 2020 |
| 82 | MnC1t3p1_G9 | 14 | IGHV3-23 | 10 | 13 | IGLV7-46 | 4 | 9 | SARS-CoV-2 convalescent - memory B cell (IgG) | Kreer et al., Cell, 2020 |
| 83 | MnC4t2p1_B3 | 14 | IGHV3-9 | 10 | 17 | IGKV1-12 | 4 | 9 | SARS-CoV-2 convalescent - memory B cell (IgG) | Kreer et al., Cell, 2020 |
| 84 | DH1210 | 14 | IGHV1-69-2 | 6 | 12 | IGLV1-47 | 8 | 12 | SARS-CoV-2 convalescent - memory B cell or plasmablast | Li et al., Cell, 2021 |
| 85 | COV2-2562 | 14 | IGHV1-8 | 8 | 16 | IGLV1-44 | 6 | 11 | SARS-CoV-2 convalescent - class switched memory B cell | Zost et al., Nature Medicine, 2020 |
| 86 | C128 | 15 | IGHV3-23 | 9 | 18 | IGKV3-20 | 6 | 10 | SARS-CoV-2 convalescent - memory B cell (IgG) | Robbiani et al., Nature, 2020 |
| 87 | COV2-2098 | 18 | IGHV3-23 | 11 | 10 | IGKV3-15 | 7 | 9 | SARS-CoV-2 convalescent - class switched memory B cell | Zost et al., Nature Medicine, 2020 |
| 88 | COV2-2308 | 18 | IGHV3-23 | 11 | 10 | IGKV3-15 | 7 | 9 | SARS-CoV-2 convalescent - class switched memory B cell | Zost et al., Nature Medicine, 2020 |
| 89 | HbnC3t1p1_F4 | 18 | IGHV3-30 | 11 | 13 | IGKV1-5 | 7 | 9 | SARS-CoV-2 convalescent - memory B cell (IgG) | Kreer et al., Cell, 2020 |
| 90 | CnC2t1p1_E8 | 20 | IGHV1-2 | 14 | 13 | IGLV2-23 | 6 | 10 | SARS-CoV-2 convalescent - memory B cell (IgG) | Kreer et al., Cell, 2020 |
| 91 | DH1073 | 21 | IGHV1-46 | 15 | 15 | IGKV3-11 | 6 | 11 | SARS-CoV-1 convalescent - memory B cell | Li et al., Cell, 2021 |
| 92 | CnC2t1p1_G6 | 21 | IGHV1-2 | 14 | 13 | IGLV2-23 | 7 | 10 | SARS-CoV-2 convalescent - memory B cell (IgG) | Kreer et al., Cell, 2020 |

**Supplementary Table 2: Fab HbnC3t1p1_C6 - PDB entry 7B0B.**

| Parameter | |
|---|---|
| Wavelength (Å) | 1.5418 |
| Space group | $P\ 2_1\ 2_1\ 2_1$ |
| Cell dimensions | |
| a, b, c (Å) | 84.41 109.10 168.17 |
| α, β , γ ° | 90 90 90 |
| Resolution (Å) | 39.23-2.98 (3.08-2.98)[a] |
| $R_{meas}$ (%) | 16.3 (64.2) [a] |
| $CC_{1/2}$ | 89.45(56.4) [a] |
| *I*/□*I* | 4.1(7.7) [a] |
| Completeness (%) | 77.27 (16.81) [a] |
| Multiplicity | 7(1.5)[a] |
| Reflections | 831358 |
| Unique reflections | 25113 (537) |
| Refinement | |
| Resolution (Å) | 39.23 – 2.98 |
| No. of reflections | 25008 (536) |
| $R_{work}/R_{free}$ (%) | 20.56 / 23.96 |
| No. of atoms | |
| Protein | 4853 |
| Ligand/ion | 28 |
| B factors | |
| Protein | 60.70 |
| Ligand/ion | |
| Ramachandran | |
| Favored (%) | 95.48 |
| Allowed (%) | 4.36 |
| Outlier (%) | 0.16 |
| Root mean square deviations | |
| Bond length (Å) | 0.009 |
| Bond angles ° | 1.32 |

[a] Values in parentheses are for the highest resolution-shell

**Supplementary Table 3: Initial and additional IGHV1-58/IGKV3-20 antibodies**

| # | Name | Total mutations (aa) | VH gene | VH mutations (aa) | CDRH3 length (aa) | VL gene | VL mutations (aa) | CDRL3 length (aa) | Origin | Reference |
|---|---|---|---|---|---|---|---|---|---|---|
| 5 | C125 | 1 | IGHV1-58 | 1 | 16 | IGKV3-20 | 0 | 9 | SARS-CoV-2 convalescent - memory B cell (IgG) | Robbiani et al., Nature, 2020 |
| 7 | HbnC3t1p1_C6 | 1 | IGHV1-58 | 0 | 16 | IGKV3-20 | 1 | 9 | SARS-CoV-2 convalescent - memory B cell (IgG) | Kreer et al., Cell, 2020 |
| 12 | COV2-2196 | 2 | IGHV1-58 | 2 | 16 | IGKV3-20 | 0 | 10 | SARS-CoV-2 convalescent - class switched memory B cell | Zost et al., Nature Medicine, 2020 |
| 32 | HbnC3t1p2_C6 | 4 | IGHV1-58 | 4 | 16 | IGKV3-20 | 0 | 9 | SARS-CoV-2 convalescent - memory B cell (IgG) | Kreer et al., Cell, 2020 |
| 43 | MnC5t2p1_G1 | 5 | IGHV1-58 | 2 | 16 | IGKV3-20 | 3 | 9 | SARS-CoV-2 convalescent - memory B cell (IgG) | Kreer et al., Cell, 2020 |
| 93 | Ab_58G6 | 7 | IGHV1-58 | 4 | 16 | IGKV3-20 | 3 | 9 | SARS-CoV-2 convalescent | Li et al., Nature Communications, 2021 |
| 94 | C043 | 17 | IGHV1-58 | 11 | 16 | IGKV3-20 | 6 | 9 | SARS-CoV-2 convalescent | Gaebler et al., Nature, 2021 |
| 95 | C598 | 7 | IGHV1-58 | 3 | 16 | IGKV3-20 | 4 | 9 | SARS-CoV-2 convalescent | Gaebler et al., Nature, 2021 |
| 96 | C827 | 12 | IGHV1-58 | 8 | 16 | IGKV3-20 | 4 | 9 | mRNA vaccinated | Wang et al., Nature, 2021 |
| 97 | CQTS004 | 6 | IGHV1-58 | 5 | 16 | IGKV3-20 | 1 | 9 | SARS-CoV-2 convalescent - memory B cell (IgG) | CN111793129A, 2020 |
| 98 | Turner-07-2C08 | 9 | IGHV1-58 | 4 | 16 | IGKV3-20 | 5 | 9 | mRNA vaccinated | Turner et al., Nature, 2021 |
| 99 | COV2-2072 | 3 | IGHV1-58 | 2 | 16 | IGKV3-20 | 1 | 9 | SARS-CoV-2 convalescent - class switched memory B cell | Zost et al., Nature Medicine, 2020 |
| 100 | P008_086 | 6 | IGHV1-58 | 2 | 16 | IGKV3-20 | 4 | 9 | SARS-CoV-2 convalescent | Graham et al., Immunity, 2021 |
| 101 | C597 | 7 | IGHV1-58 | 4 | 16 | IGKV3-20 | 3 | 9 | SARS-CoV-2 convalescent | Gaebler et al., Nature, 2021 |
| 102 | COV2-3025 | 4 | IGHV1-58 | 4 | 16 | IGKV3-20 | 0 | 10 | SARS-CoV-2 convalescent - class switched memory B cell | Zost et al., Nature Medicine, 2020 |
| 103 | I14 | 7 | IGHV1-58 | 4 | 16 | IGKV3-20 | 3 | 9 | SARS-CoV-2 convalescent | Adreano et al., Cell, 2021 |
| 104 | P008_081 | 7 | IGHV1-58 | 2 | 16 | IGKV3-20 | 5 | 10 | SARS-CoV-2 convalescent | Graham et al., Immunity, 2021 |
| 105 | Wang-C387 | 1 | IGHV1-58 | 1 | 16 | IGKV3-20 | 0 | 9 | SARS-CoV-2 convalescent | Wang et al., Science, 2020 |
| 106 | CV07-287 | 1 | IGHV1-58 | 1 | 16 | IGKV3-20 | 0 | 9 | SARS-CoV-2 convalescent - memory B cell (IgG) | Kreye et al., Cell, 2020 |
| 107 | COV2-2381 | 4 | IGHV1-58 | 2 | 16 | IGKV3-20 | 2 | 10 | SARS-CoV-2 convalescent - class switched memory B cell | Zost et al., Nature Medicine, 2020 |
| 108 | R200-1B9 | 9 | IGHV1-58 | 6 | 16 | IGKV3-20 | 3 | 9 | SARS-CoV-2 convalescent - memory B cell (IgG) | Vanshylla et al., Cell Host & Microbe, 2022 |
| 109 | R259-1B9 | 13 | IGHV1-58 | 8 | 16 | IGKV3-20 | 5 | 9 | SARS-CoV-2 convalescent - memory B cell (IgG) | Vanshylla et al., Cell Host & Microbe, 2022 |
